# The Alzheimer’s disease-associated rs28834970 variant in *PTK2B* impairs electrical activity and regulates Tau phosphorylation and APP processing in human neuronal cultures

**DOI:** 10.64898/2026.10.26.761842

**Authors:** Nina Lannette-Weimann, Marlène Huyvaert, Floriane Limoge, Morgane Lemaire, Vance Gao, Florence Maurier, Karine Guyot, Charlotte Henocq, Audrey Coulon, Valérie Buiche, Florian Rabiller, Nad’a Majerníková, Pauline Walle, Sophie Desnoulez-Salomé, Bruno Neiva-Queirós, Pol López-Pérez, Leo Morita Miyakoshi, Ana Raquel Melo de Farias, Marcos R. Costa, Philippe Amouyel, Chloé Najdek, Angelika Dannert, Judit González-Gallego, Manon Muntaner, Benjamin Grenier-Boley, Julien Chapuis, Dominik Paquet, Devrim Kilinc, Jean-Charles Lambert, Julie Dumont

## Abstract

**Background:** Genome-wide association studies have identified the *PTK2B* gene, which encodes Proline-rich tyrosine kinase 2 (PYK2), as a genetic risk factor for Alzheimer’s disease (AD). However, its role in AD aetiology remains to be understood, as a complex contribution to amyloid β (Aβ) toxicity, tauopathy and neuronal dysfunction is suggested by contradictory evidence. Determining how the causal variant among AD-associated *PTK2B* variants impairs PYK2 function to trigger AD hallmarks could bring clarification.

**Methods:** We combined *in silico* prediction tools with luciferase reporter assays in neuroblastoma cells to prioritize potential AD-associated *PTK2B* functional variants. We generated isogenic KOLF2.1J induced pluripotent stem cells (iPSCs) carrying the most relevant functional variant using CRISPR-Cas9 and differentiated them into human neuronal cultures (hNCs). We assessed PYK2 expression via immunoblotting, performed transcriptomic profiling using RNA-sequencing, analyzed neuronal network activity *via* multielectrode array (MEA) recordings, quantified synaptic density through immunocytochemistry in microfluidic devices, and assessed Tau phosphorylation and amyloid-precursor protein (APP) metabolism.

**Results:** The rs28834970 regulatory variant was identified as the most functionally relevant in our model, with the C risk allele promoting transcription factor C/EBPβ binding within a cis-regulatory element. In hNCs, the homozygous rs28834970 CC genotype reduced total PYK2 levels by more than 80%, while increasing the proportion of phosphorylated (activated) PYK2. Transcriptomic analysis revealed 724 downregulated and 113 upregulated gene sets, with altered expression of genes involved in extracellular matrix organization, cytoplasmic translation and neurotransmitter secretion. MEA recordings showed reduced network recruitment and altered burst organization. Immunocytochemistry in microfluidic devices consistently revealed that rs28834970 impacts dendritic development and postsynaptic maturation. The risk genotype significantly elevated Tau phosphorylation at pathological sites (Thr181, Thr217, Thr231, Ser202-Thr205, Ser396-Ser404), and enhanced APP processing, resulting in decreased levels of APP and CTF but increased levels of sAPPα, sAPPβ and Aβ.

**Conclusion:** The rs28834970 C risk allele mechanistically links AD susceptibility to downregulation of *PTK2B*/PYK2 expression and to the increase in Tau pathology and Aβ levels while compromising synaptic structure and function.

## INTRODUCTION

The three main pathological hallmarks of Alzheimer’s disease (AD) are (i) senile plaques, which are composed of accumulated oligomeric amyloid β (Aβ) peptides in the extracellular space, (ii) neurofibrillary tangles, which are formed by aggregated, abnormally phosphorylated Tau proteins within neurons, and (iii) early synaptic dysfunction and loss. However, the initial events that trigger the early synaptic dysfunction and lesion development observed in AD patients (and in AD-like models), remain unclear.

Large twin studies have shown that the heritability of AD is particularly high [1], making the delineation of its genetic architecture a relevant tool for understanding the underlying pathophysiological processes. To characterize this AD genetic component, several genome-wide association studies (GWAS) have been conducted over the last 15 years to identify genes modulating AD risk, and a recent GWAS meta-analysis confirmed 56 loci specifically associated with clinically diagnosed AD, including *PTK2B* [2,3]. Yet, like many GWAS-identified genes, the pathophysiological role of *PTK2B* in AD and the functional consequences of its associated variants remain poorly characterized.

*PTK2B* encodes PYK2, a Ca²⁺-activated non-receptor tyrosine kinase that belongs to the focal adhesion kinase (FAK) family. PYK2 mediates cell-extracellular matrix interactions and transduces extracellular signals regulating cell migration and survival. Although *PTK2B* is broadly expressed throughout the brain (**Supplementary fig. 1a-c**), it is particularly enriched in hippocampal neurons [4]. In the middle temporal gyrus of AD patients, *PTK2B* exhibits heterogeneous expression patterns across disease progression, depending on cell type. (**Supplementary fig. 1 d-g**) [5,6]. At the protein level, PYK2 activation is tightly coupled to intracellular calcium and neuronal activity and excitatory neurotransmission [7,8]. Calcium/calmodulin binding is thought to facilitate PYK2 dimerization, enabling intermolecular autophosphorylation at Tyr402 [9,10]. Once activated, PYK2 recruits numerous binding partners, including Src-family kinases, which reciprocally activate PYK2. Through distinct phosphorylated sites and protein interactions, PYK2 participates in diverse neuronal signaling processes [11]. Under physiological conditions, PYK2 supports neurite formation and maintenance, synaptic plasticity, and NMDA receptor (NMDAR) signaling [11]. Studies using PYK2 knockout mice show that hippocampal PYK2 is essential for spatial memory and long-term potentiation (LTP), and regulates dendritic spine density, morphology, and postsynaptic organization [12]. In the postsynaptic compartment, PYK2 interacts with PSD-95 and phosphorylates NMDAR subunits (GluN2A/B) directly or through the activation of SRC, which is also an AD genetic risk factor [3]. These findings highlight the critical role of PYK2 in synaptic plasticity, a process central to cognitive decline in AD.

The role of PYK2 in AD pathology is controversial and context-dependent. Regarding the amyloid component, Salazar et al. showed that *Pyk2* deletion protects hippocampal slices from Aβ-mediated suppression of LTP and that *Pyk2* is required for synaptic marker loss and memory impairment in APPswe/PSEN1ΔE9 mice [13]. This finding was extended by demonstrating that genetic deletion of *Pyk2* prevents Aβ-induced hippocampal neuronal hyperactivity and dendritic spine/synaptic loss [14], which are early markers of AD [15,16]. However, in contrast to these observations, it has been shown that Pyk2 deletion did not improve the phenotype of 5xFAD mice [17]. Conversely, Pyk2 overexpression in this model improved behavioral and neuropathological outcomes. Model-dependent divergences extend to tauopathy. *PTK2B* was initially described as a modulator of Tau toxicity in a *Drosophila* model, interacting *in vitro* with Tau and co-localizing with hyperphosphorylated and oligomeric Tau in AD brains [18]. While Tau pathology was enhanced by *Pyk2* overexpression in Tau transgenic pR5 mice overexpressing human full-length P301L Tau [19], *Pyk2* deletion increased Tau phosphorylation and exacerbated Tau-associated phenotypes in PS19 transgenic mice [20]. These model-dependent divergences raise critical questions about the potential benefits of PYK2 inhibition or activation in AD. Elucidating the role of AD-associated *PTK2B* variants in the aetiology of AD may address this question. In this study, we aimed to identify functional *PTK2B* variants and characterize their impact on PYK2 and AD hallmarks in a human induced pluripotent stem cell (iPSC)-derived neuronal model.

## METHODS

### *In silico* functional prediction of AD-associated *PTK2B* variants

Functional annotation of AD-associated *PTK2B* gene variants was performed using the UCSC Genome Browser [21] to integrate transcript, regulatory, and epigenomic information. Variants were annotated according to the MANE (Matched Annotation from NCBI and EMBL-EBI) transcript set to ensure standardized transcript assignment [22]. Regulatory annotations were obtained from the ENCODE4 Registry of candidate cis-Regulatory Elements (cCREs), while chromatin accessibility and epigenomic features were assessed using ENCODE datasets, including DNase-seq and ATAC-seq signal tracks, histone modification tracks (H3K4me3 and H3K27ac), and CTCF binding profiles [23,24]. To maximize tissue relevance, ENCODE datasets were restricted to experiments performed in human brain tissues and brain primary cells. Transcription factor binding data were obtained from the ReMap ChIP-seq database [25]. ATAC-seq data in neurons and non-neuronal cells across 14 distinct human brain regions were obtained from the online database “Brain Open Chromatin Atlas” (BOCA) [26]. Sequence-based functional predictions for AD-associated *PTK2B* gene variants were generated using the Sei model, derived from the DeepSEA framework [26], and the maximum sequence class score was extracted for each variant as a measure of predicted regulatory activity. This score calculates the computational difference in chromatin profiles between a reference allele and an alternative mutated allele. Sei algorithm [26] was also used to predict whether a sequence variation will disrupt or create a binding site for specific transcription factors. Last, Position Weight Matrix (PWM) from the JASPAR 2026 database [27] was extracted for C/EBPβ to illustrate the potential impact of rs28834970 on C/EBPβ binding within *PTK2B* intron 6.

### Analysis of single-nucleus RNA-sequencing datasets of human brains

To assess *PTK2B* expression according to the cell type, we analyzed two publicly available single-nucleus RNA-seq datasets of human brain tissue: Li *et al.,* 2025 [28] performed on temporal cortex tissue (GSE237718) and Leng *et al*., 2021 [29] performed on entorhinal cortex and superior frontal gyrus (syn21788402). Analysis was conducted using the Seurat R package [30]. For the Li *et al*., 2025 [28] dataset, cells were filtered to retain ≥200 detected features and <10% mitochondrial genes, with features detected in ≥10 cells. Following SCTransform normalization with mitochondrial gene regression [31], FindNeighbors was applied using 30 principal component analysis (PCA) dimensions, and FindClusters with resolution 0.35. Cell types were classified using marker genes: microglia (PTPRC), astrocytes (SLC1A2), oligodendrocytes (PLP1), oligodendrocyte precursor cells (VCAN), inhibitory neurons (GAD1, excluding VCAN+ clusters), excitatory neurons (CAMK2A, excluding GAD1+ clusters), and vascular/endothelial cells (CLDN5). For the Leng *et al*., 2021 dataset [29], cell types were pre-annotated.

To investigate whether *PTK2B* expression varies with the progression of AD, we analyzed the SEA-AD datasets for the middle temporal gyrus (MTG) [5,6] in which the Continuous Pseudoprogression Score (CPS) deriving from quantitative neuropathology orders donors along a continuous AD pathological burden gradient.

### Cell culture

#### SH-SY5Y neuroblastoma cells

The SH-SY5Y neuroblastoma cell line was purchased from the American Tissue Culture Collection (ATCC, CRL-2266). Cells were grown in a 1:1 mixture of Eagle’s Minimum Essential Medium (ATCC® 30-2003) and F-12 Medium (Gibco®), supplemented with 10% heat-inactivated Fetal Bovine Serum (FBS), 2mM L-Glutamine and 100U/mL antibiotics (penicilline and streptomycine). Cell cultures were maintained at 37°C with 5% CO_2_. All experiments were performed using cells in passages 5 to 10.

#### IPSC maintenance

KOLF2.1J iPSCs [32] (Applied StemCell, CA, USA) were maintained at 37°C in a humidified atmosphere containing 5% CO₂. Cells were cultured in E8 Flex medium (A2858501, Gibco, Thermo Fisher Scientific), which was fully replaced at least every 3 days. Cells were split before colonies became too dense or exceeded 80% confluency at a 1:6 ratio using non-enzymatic dissociation with 0.5 mM EDTA for 5 min at room temperature, allowing cells to remain in small colonies. Cells were seeded onto plates pre-coated with vitronectin (0.5 µg/cm²; A14700, Gibco, Thermo Fisher Scientific).

#### Human iPSC-derived neuronal cultures

NPCs were generated from KOLF2.1J iPSCs using the STEMdiff SMADi Neural Induction kit (08582, STEMCELL Technologies), according to the manufacturer’s instructions with minor modifications. Briefly, 2.5 × 10⁶ iPSCs were seeded per well at D0 and maintained in Neural Induction Medium supplemented with SMADi for 21 days. Cells were passaged at D7 (2.5 × 10⁶ cells/well) and D14 (2.7 × 10⁶ cells/well). At D21, NPCs were harvested and cryopreserved at 4 × 10⁶ cells/vial and efficiency of neural induction was assessed by immunocytochemistry of PAX6, SOX1, Nestin, OCT4 and SOX10 markers. iPSCs used for neural induction were within ten passages of the cultures used for quality control, and pluripotency was confirmed by immunofluorescence before each induction. Five independent neural inductions were performed in 6-well plates pre-coated with 15 µg/mL poly-L-ornithine (PLO; P4957 Sigma) and 10µg/mL Laminin (Lam, L2020, Sigma), with both genotypes processed in parallel in each induction. Neural progenitor cells (NPCs) were maintained and expanded in STEMdiff Neural Progenitor Medium (NPM; 05833, STEMCELL Technologies) prior to differentiation and were used at a maximum of passage two.

Five independent neuronal differentiation experiments were performed, with both genotypes processed in parallel in each experiment. For standard differentiation, NPCs were seeded in PLO/Lam coated 24-well plate at 1 × 10⁵ cells/well in 500 µL NPM supplemented with 10 µM Y- 27632 ROCK inhibitor (T1725, Target Mol). At D1, 500 µL BrainPhys Neuronal Medium supplemented with 2% NeuroCult SM1 Neuronal Supplement, 1% N2 Supplement-A, 20 ng/mL BDNF, 20 ng/mL GDNF (BrainPhys hPSC Neuron Kit, 05795, STEMCELL Technologies), 1 mM dibutyryl-cAMP (73886, STEMCELL Technologies), 200 nM ascorbic acid (105021000, Thermo Scientific) and 1 µg/ml Laminin. Half-medium changes with this BrainPhys differentiation medium were performed at D3, D7, D10, D14, D17, D21, and D24, and cultures were maintained until D28.

### Luciferase reporter assays

Luciferase reporter vectors containing *PTK2B* gene sequence with either the common allele or the minor allele for the seven AD-associated variants were obtained from genomic DNA of individuals with known contrasted genotypes. They were generated by subcloning the PCR product amplified using oligonucleotides containing KpnI/NheI restriction sites (**Supplementary table 4**) upstream the SV40 promoter and firefly luciferase gene in the pGL3-promoter vector (Promega). The insert identity was verified by Sanger sequencing (Genoscreen). Plasmids were purified using NucleoBond® Xtra Midi/Maxi kit (Macherey-Nagel) and quantified using the NanoVue Plus spectrophotometer (Biochrom).

SH-SY5Y cells were plated in 24-well plates at a seeding density of 100,000 cells/cm^2^. When cells reached 70-80% confluence, they were transfected with 200 ng *Firefly* luciferase reporter vector. The *Renilla* luciferase expression vector (pRL, Promega, 100 ng) was used for normalization. Transfections were carried out in 100µL OptiMEM^®^ (Gibco) using FuGENE^®^ HD (Promega) transfection reagent with a DNA (μg): FuGene HD (μL) ratio of 1:3 and according to the manufacturer’s instructions. 48 h after transfection, *Firefly* and *Renilla* luciferase assays were performed using the Dual-Luciferase Reporter Assay System kit (Promega) according to the manufacturer’s instructions. Luminescence was recorded with a Wallac-Victor luminometer (Perkin-Elmer).

### CRISPR-CasG editing of KOLF2.1J iPSCs and quality control

Guide RNA and single-stranded oligodeoxynucleotide (ssODN) repair template sequences are provided in **Supplementary table 4**. CRISPR-Cas9 editing in KOLF2.1J iPSCs [32] was achieved by nucleofecting ribonucleoprotein complex formed by mixing 60 pmol of guide RNA with 30 pmol of Sp. Cas9 protein at room temperature for 20 min. Just before nucleofection, we added 120 pmol of repair template designed to introduce the C allele through homology-directed repair. iPSCs were detached with Accutase (A6924, Sigma), dissociated into a single-cell suspension, and 250,000 cells were resuspended in P3 buffer (Primary Cell 4D-Nucleofector X Kit, Lonza) before being nucleofected with the RNP and the repair template using the Lonza 4D-Nucleofector program CA137. Individual colonies were picked and isolated in 96-well plates screened for the introduction of the homozygous C allele using restriction fragment length polymorphism analysis with HpyCH4V restriction enzyme (New England Biolabs). Sanger sequencing verified the intended T>C homozygous substitution for the rs28834970 variant. Quality control of the correctly edited clones was performed according to Weisheit *et al*., 2021 [33]. CRISPR-edited iPSCs retained typical morphology and pluripotency markers (SOX2, OCT4, SSEA4) by brightfield microscopy and immunofluorescence. Genomic DNA from edited and parental KOLF2.1J iPSCs was extracted and fingerprinted at the D1S80 locus (chromosome 1) using PCR and agarose gel analysis to confirm cell identity. CRISPR-induced alterations were assessed via quantitative genomic PCR using TERT as reference and whole-genome sequencing (WGS), with karyotypic integrity confirmed by qPCR (hPSC Genetic Analysis Kit, StemCell technologies) and WGS (Beijing Genomics Institute). WGS was analyzed as previously described in details[34], using the nf-core sarek (v3.5.1) pipeline[35] with fastp (v0.24.0), dragmap (v1.2.1), and Picard MarkDuplicates (v3.3.0). Variant calling used GATK HaplotypeCaller (v4.6.1.0), chromosome copy number was estimated via Tiddit (v6.6.1), and variants were annotated using Ensembl’s VEP (v111). Structural variants (SVs) were detected with DELLY [36] (v1.7.2), Manta [37] (v1.6.0), and Smoove (v0.2.8; https://github.com/brentp/smoove) coupled with Duphold (v0.2.3) [38]. SVs from each tool were then merged using SURVIVOR [39] (v1.0.7, with 500 bp breakpoint distance). Only variants identified by all three tools with consistent genotypes were retained; sample contamination was assessed with VerifyBamID (v2.0.1). WGS confirmed the homozygous T>C substitution at chr8: 27337604 (Hg38). No additional variants were detected in the *PTK2B* region. Quality control confirmed normal ploidy and no CRISPR-induced large deletions, insertions, rearrangements, or loss of heterozygosity. No off-target variants were detected at the 209 predicted CRISPOR [40] sites (**Supplementary table 5**).

### Microfluidic device fabrication and neuronal differentiation therein

Masters of microfluidic devices were fabricated through photolithography as previously described [41]. Devices were cast from polydimethylsiloxane (PDMS, Sylgard 184, Dow Corning, Midland, MI) and irreversibly bonded in pairs to glass coverslips using an O_2_ plasma generator (Diener, Ebhausen, Germany). Prior to cell culture, devices were sterilized under UV (Light Progress, Anghiari, Italy) for 20 min and coated with 20 µg/mL poly-L-lysine (PLL, Sigma, catalog #P6282) diluted in borate buffer (0.31% boric acid, 0.475% sodium tetraborate, pH = 8.5) followed by 20 µg/mL laminin (CC095, Sigma-Aldrich) diluted in 0.01 M PBS with Ca^2+^ and Mg^2+^ (Gibco, ThermoFisher Scientific). For synapse imaging and quantitative analysis, tricompartment microfluidic devices that isolate the neuronal network from cell bodies were used [42] (Fig. 5a). NPCs^WT^ and NPCs^HOM^ were seeded in mirror in two microfluidic devices bounded on the same coverslip side by side. Three independent cultures were performed. At D0, 6–12 × 10³ NPCs were plated in presynaptic and postsynaptic chambers in NPM supplemented with 10 µM Y-27632 ROCK inhibitor. Neural differentiation was induced at D1 by half-medium change with complete BrainPhys™ Neuronal Medium and hNCs were maintained for four weeks with half medium changes twice a week.

### Immunocytochemistry

Cells were fixed with 4% paraformaldehyde (EMS, Hatfield, PA) for 15 min at RT, permeabilized with 0.3% Triton X-100 in PBS for 5 min at RT, and blocked with 3% BSA and 0.1% Triton X-100 for 1 h at RT. After blocking, cells were incubated overnight at 4°C with the following primary antibodies: rat anti-SOX2 (Invitrogen, catalog #14-9811-82, 1:100 dilution), rabbit anti-OCT4 (Invitrogen, catalog #710788, 1:250 dilution), mouse anti-SSEA4 (StemCell Technology, catalog #60062, 1:200 dilution), rabbit anti-PAX6 (BioLegend, catalog #901301, 1:50 dilution), mouse anti-Nestin (StemCell Technology, catalog #60091, 1:1000 dilution), goat anti-SOX1 (RCD Systems, catalog #AF3369, 1:200 dilution), chicken anti-MAP2 (SySy antibodies, catalog #188006, 1:500 dilution), rabbit anti-GFAP (Merck, catalog #AB5804, 1:1000 dilution), and mouse anti-PYK2 (Cell Signaling, catalog #3480, 1:100 dilution). After washes with PBS, cells were incubated for 2 h at RT with the following secondary antibodies (1:400 dilution): Donkey anti-Chicken Alexa 488 (Jackson Immunoresearch, 703-545-155), Donkey anti-Rabbit Alexa 594 or 647 (Jackson Immunoresearch, 711-585-152, and 711-605-152), Donkey anti-Mouse Alexa 594 or 647 (Jackson Immunoresearch, 715-585-151, and 715-605-151), Donkey anti-Goat Alexa 488 (Jackson Immunoresearch, 705-545-147), Donkey anti-Rat Alexa 488 (Jackson Immunoresearch, 712-545-153).

hNCs in microfluidic devices were fixed in 4% paraformaldehyde in PBS for 15 min at RT, washed three times with PBS, and permeabilized with 0.3% Triton X-100 in PBS for 5 min at RT. Cells were blocked in PBS containing 5% normal donkey serum for 3 h at RT before overnight incubation at 4°C with the following primary antibodies: chicken anti-MAP2 (SySy antibodies, 188006, 1:50 dilution), guinea pig anti-Homer (SySy antibodies, 160318, 1:200 dilution), and mouse anti-Synaptophysin (SySy antibodies, 101011, 1:100 dilution). Cells were washed twice with PBS and incubated for 2 h at RT with the following secondary antibodies (1:500 to 1:250 dilution): Donkey anti-Chicken DyLight™ 405 (Jackson Immunoresearch, 703-475-155), Donkey anti-Guinea Pig AlexaFluor 594 (Jackson Immunoresearch, 706-585-148), Donkey anti-Mouse AlexaFluor 488 (Jackson Immunoresearch, 715-545-151). Cells were rinsed three times with PBS and access wells were filled with 90% glycerol.

### Synaptic marker imaging and analysis

Synaptic compartments of microfluidic chips were imaged *via* XLight V3 spinning disk confocal microscope (Nikon) using a 60× objective and a *z*-stack interval of 0.2 µm. Typically, ten 1024×1024-pixel images were acquired per synaptic chamber. Images were deconvoluted using the 3D deconvolution module of the Nikon Imaging Software with default parameters. Deconvoluted images were analyzed using Imaris V11 (Oxford Instruments) by segmentation of the MAP2 signals via the Surfaces module and of Synaptophysin and Homer1 signals via the Spots module. Spots were classified according to the shortest distance to the complementary synaptic spot population, i.e., Synaptophysin spots were classified according to their distance to Homer1 spots, and vice versa. Distance-based classification was performed using the 1D Classification tool, and six classes were defined: <0.2 µm, 0.2-0.3 µm, 0.3-0.4 µm, 0.4-0.5 µm, 0.5-0.6 µm, >0.6 µm. This classification enabled the quantification of the spatial relationship between pre- and postsynaptic puncta both in the entire image. Surface and spot data were exported from Imaris and analyzed via a custom Matlab (MathWorks) code that calculates (for each image) the total volume of MAP2 surfaces, the numbers of Synaptophysin and Homer1 spots, and the fraction of Synaptophysin spots assigned (%SYP_assigned), i.e., in the proximity of a Homer1 spot, as a function of the distance cut-off. The distance cut-off was empirically determined separately for each of the three independent experiments to ensure that it is on linear portion of the %SYP_assigned vs. distance cut-off curve [42]. This resulted in distance cut-offs of 0.4 µm for one experiment and 0.5 µm for the remaining two experiments.

### Immunoblotting

Protein quantification was performed using the BCA protein assay (ThermoFisher Scientific). Equal amounts (5–20 μg) of cell lysate were collected in RIPA buffer (0.05 M Tris, 0.15 M NaCl, 1% NP-40, 0.1% SDS, 1 mM sodium orthovanadate, and 0.5% sodium deoxycholate, pH = 7.4) containing protease inhibitors (Complete mini, Roche Applied Science, Penzberg, Germany), lithium dodecyl sulfate (ThermoFischer Scientific), and reducing agent (Invitrogen). Samples were sonicated at 60% and denaturated at 95°C for 5 min. Protein extracts were separated in 4- 12% Bis-Tris, or 3-8 % Tris-Acetate SDS–polyacrylamide gels (NuPAGE, Thermo Scientific) and transferred on to nitrocellulose membranes (Bio-Rad). Membranes were blocked in SuperBlock (ThermoFisher Scientific) for 1 h at RT and incubated with primary antibodies at 4°C overnight. The following primary antibodies were used: mouse anti-β-Actin (Sigma Aldrich, catalog #A1978, 1:10,000 dilution), mouse monoclonal anti-PYK2 (Cell Signaling, catalog #3480, 1:1000 dilution) and rabbit polyclonal anti-PYK2 (Sigma Aldrich, catalog #P3902, 1:1000 dilution), rabbit anti-phospho-PYK2 Tyr402 (Cell Signaling, catalog #3291, 1:1000 dilution), rabbit anti-ST8SIA5 (Thermo Fisher, catalog#PA5112727, 1:1000 dilution), rabbit anti-NRXN3 (Abcam, catalog#ab192313, 1:1000 dilution), rabbit anti-total Tau (Dako, catalog #A0024, 1:10,000 dilution), mouse anti-phospho-Tau Thr181 (D9F4G) (Cell Signaling, catalog #12885, 1:1000 dilution), rabbit anti-phospho-Tau Thr217 (Thermo-Fisher, catalog #44-744, 1:1000 dilution), mouse anti-phospho-Tau Thr231 (Thermo-Fisher, catalog #MN1040, 1:500 dilution), mouse anti-phospho-Tau Ser202/Thr205 (AT8) (Thermo Fisher Scientific/Invitrogen, catalog #MN1020, 1:500 dilution), mouse anti-phospho-Tau Ser396/Ser404 (PHF1) (Peter Davies, 1:1000 dilution), rabbit anti-APP C-term (CTF) (Sigma Aldrich, catalog #A8717, 1:5000 dilution), guinea pig anti-APP C-term (total APP) (SySy antibodies, catalog #127005, 1:1000 dilution), mouse anti-β-Amyloid 1-16 (clone 6E10, sAPPα) (Biolegend, catalog #803015, 1:1000 dilution), rabbit anti-sAPPβ (Biolegend, catalog #813401, 1:1000 dilution), rabbit anti-p38 MAPK and anti-phospho-p38 MAPK Thr180/Tyr182 (Cell Signaling, catalog #9212 and #4511, 1:1000 dilution), rabbit anti-ERK1/2 (Abcam, catalog #ab17942, 1:1000 dilution), rabbit anti-phospho-ERK1/2 Thr202/Tyr204 (RCD Systems, catalog #MAB1018, 1:1000 dilution), mouse anti-GSK3β and anti-phospho-GSK3β Tyr216 (BD Transduction Laboratories, catalog #610202 and #612313, 1:5000 and 1:1000 dilution, respectively), rabbit anti-AKT and anti-phospho-AKT Ser473 (Cell Signaling, catalog #4691 and #4060, 1:1000 dilution). The following day, membranes were incubated with HRP-conjugated secondary antibodies for 2 h at RT and revealed using Immobilon Classico Western HRP Substrat (Millipore) and Amersham Imager 600 (GE Life Sciences). Secondary antibodies (1:5000 dilution) used were Goat anti-Chicken HRP (ThermoFisher Scientific, A16054), Goat anti-Guinea pig HRP (Jackson Immunoresearch, catalog #106-035-003), Goat anti-Rabbit HRP (Jackson Immunoresearch, catalog #111-035-003) and Goat anti-Mouse HRP (Jackson Immunoresearch, catalog #115-035-003). Chemiluminescence was quantified using the Amersham Imager 600 (GE Healthcare) imaging system. Densitometry of bands was quantified using ImageJ (v2.14) [43].

### Alpha-LISA measurements

Conditioned media from hNCs were collected at 4 weeks of differentiation to dose Aβ_1-x_ using Alpha-LISA assays (Alpha-LISA Amyloid β1–X Kit, AL288C, PerkinElmer) according to the manufacturer’s instructions. 5 μL of cell medium or standard solution was added to an Optiplate-96 microplate (PerkinElmer), incubated with 5 μL of 10X mixture of acceptor beads and biotinylated antibody for 1 h at RT, and then by 40 μL of 1.25X donor beads for 30 min at RT. Luminescence was measured using an EnVision-Alpha Reader (PerkinElmer) at 680 nm excitation and 615 nm emission wavelengths. Aβ concentrations were normalized by the quantity of protein measured using BCA Protein Assay (Thermo Scientific, 23225).

### Microelectrode array

The MEA recordings were carried out using 48-well CytoView MEA plates (M768-tMEA-48W, AXION Biosystems), with each well containing 16 electrodes. MEA plates were pre-coated with 10 µg/mL poly-L-lysine (P6282, Merck) and 20 µg/mL laminin (23017015, Gibco, Thermo Fisher Scientific). NPCs were seeded at 3.5 × 10⁴ cells/well in a 5 µL droplet of NPM supplemented with 10 µM Y-27632 ROCK inhibitor. After 45 min, 150 µL of the same medium was added to each well. At DIV1, 150 µL of BrainPhys neuronal differentiation medium was added, and subsequent half-medium changes were performed. Extracellular action potentials were recorded each week until DIV28 using MaestroPro (Axion Biosystems). Five-minute-long recordings from three independent cultures were analyzed for multi-unit activity using a 6 standard deviation adaptive threshold to detect spikes, with bursts defined as ≥5 spikes within 100 ms.

### Bulk RNA sequencing and data processing

Total RNA extraction was performed using the NucleoSpin RNA Mini kit (Macherey Nagel, 740955.50). Library preparation and sequencing were outsourced to Novogene Europe (Planegg, Germany). Libraries were sequenced on the NovaSeq X Plus Series platform (PE150) with a target of 18 Gb of raw data per sample.

Raw sequencing data were processed using the nf-core/rnaseq workflow (version 3.26.0) (https://doi.org/10.5281/zenodo.20072251), executed with the default pipeline parameters. The default STAR–Salmon processing route was used, consisting of splice-aware read alignment with STAR [44], followed by transcript-level quantification with Salmon [45]. Sequencing data were analyzed as unstranded paired-end reads of 150 bp. Reads were aligned to the human reference genome GRCh38.p12, corresponding to NCBI assembly accession GCA_000001405.27. Gene-level count data were obtained from the merged Salmon gene-count matrix (salmon.merged.gene_counts.tsv) and used as input for downstream differential gene expression analysis.

### Differential gene expression analysis

Differential gene expression analysis was performed in R using the Bioconductor package DESeq2 (version 1.52.0) [46]. Gene-level count data were obtained from the merged Salmon gene-count matrix. Genes with insufficient expression were filtered prior to statistical testing by retaining genes with a cumulative count of at least 15 across all samples from both experimental conditions. The experimental design comprised two conditions, hNCs^HOM^ and hNCs^WT^, with five biological replicates per condition. hNCs^WT^ was defined as the reference condition. The DESeq2 design formula was ∼ condition, and the differential expression analysis was performed using the hNCs^HOM^ versus hNCs^WT^ contrast. Library size normalization was performed using the median-of-ratios method implemented in DESeq2. Differential expression was assessed using a negative binomial generalized linear model, with gene-specific dispersion parameters estimated from the count data. Wald tests were used to evaluate the statistical significance of the hNCs^HOM^ versus hNCs^WT^ contrast. For each gene, the estimated log2 fold-change, standard error, Wald test statistic, nominal p-value, and Benjamini–Hochberg-adjusted p-value were obtained.

The significance threshold for differential gene expression was 0.05 for the adjusted p-value (false discovery rate, FDR). The sign of the log2 fold change was interpreted relative to the hNCs^HOM^ versus hNCs^WT^ contrast, with positive values indicating higher expression in hNCs^HOM^ and negative values indicating lower expression in hNCs^HOM^.

### Functional enrichment analysis

Functional enrichment analyses were performed using clusterProfiler version 4.20.0 [47]. Gene identifiers were converted from Ensembl gene identifiers to the Entrez Gene identifiers using the annotation information provided by org.Hs.eg.db version 3.23.1.

GSEA were performed using genes ranked according to the Wald test statistic obtained from the DESeq2 differential expression analysis. Genes for which the test statistic was not available were excluded from the ranked gene list prior to enrichment analysis. Separate GSEA were performed for different functional annotation databases, including Gene Ontology (GO) Biological Process, GO Cellular Component, GO Molecular Function, Reactome Pathways, and KEGG pathways. For all annotation collections, gene sets containing between 50 and 500 genes were retained for analysis. This uniform minimum gene-set size of 50 genes was applied across all annotation collections to reduce the influence of very small gene-sets, for which enrichment scores may be driven by a limited number of genes, and to focus the analysis on gene-sets with sufficient representation in the dataset. This criterion was also intended to provide a consistent basis for comparison of enrichment patterns across the different functional annotation databases. The maximum gene-set size was set to 500 genes to limit the contribution of very broad gene sets and to facilitate the interpretation of biologically more specific enrichment signals. For all enrichment analyses, the p-value cutoff was set to 0.05 and the q-value cutoff was set to 0.05. Multiple-testing correction was performed using the Benjamini–Hochberg procedure.

### Statistical analyses

No *a priori* statistical power calculation was performed to determine sample size. Experiments were not randomized, and investigators were not blinded to group allocation during data collection or outcome assessment, except for the generation of bulk RNA-seq data. Outliers were identified and excluded only for synaptic quantification in microfluidic device cultures, using the interquartile range method (IQR = Quartile 3 – Quartile 1), i.e., data points falling outside of the interval defined from Q1−1.5×IQR to Q3 + 1.5×IQR. Specifically, outlier images were detected per condition per experiment in terms of MAP2 volume and excluded from the analysis. Outliers in Synaptophysin spot density (spot number per MAP2 volume), Homer spot density, and synapse density were then detected and excluded from the analysis. Device means were then calculated and normalized to the mean of the control condition, separately for each independent experiment. Normalized device means were then pooled, and outlier devices were determined according to the 1.5×IQR method. Statistical tests were performed after excluding one outlier device. No other data were excluded from analysis. Longitudinal microelectrode array (MEA) recordings were analyzed using a mixed-effects model with Holm-Šídák’s multiple comparisons test. All other comparisons were performed using the Mann-Whitney U test (two-sided, exact p-values). All statistical analyses were conducted using GraphPad Prism version 11.0 (GraphPad Software, San Diego, CA, USA).

## RESULTS

### *In silico* and *in vitro* analyses of the seven AD-associated *PTK2B* variants prioritized rs28834970

GWAS meta-analyses identified seven variants within intron 6 of PTK2B whose minor alleles confer increased AD risk (**Supplementary table 1**) [2]. Although Quantitative Trait Loci eQTL associations could not be analyzed in the snRNA-seq datasets to assess whether these variants explain the *PTK2B* expression changes with disease progression (**Supplementary fig. 1 d-g**) [5,6], it was reasonable to hypothesize that one or more of these common variants might functionally modulate *PTK2B* brain expression. However, all seven variants display similar minor allele frequencies (37–41% in European populations) and cluster within a single linkage disequilibrium block (r² > 0.8), which precludes distinguishing the causal variant through statistical association alone. To identify the most functionally relevant variant, we integrated *in silico* and *in vitro* approaches in SH-SY5Y neuroblastoma cells.

We first evaluated the impact of all seven variants on gene expression using luciferase reporter assays. Vectors expressing a ∼500 bp region bearing either the common or risk allele of each variant were cloned upstream of an SV40 promoter driving luciferase expression, then transfected into SH-SY5Y cells. Only three variants significantly affected luciferase activity: rs28834970, rs6987305, and rs755951 (**Fig. 1a**).

**Figure 1.**
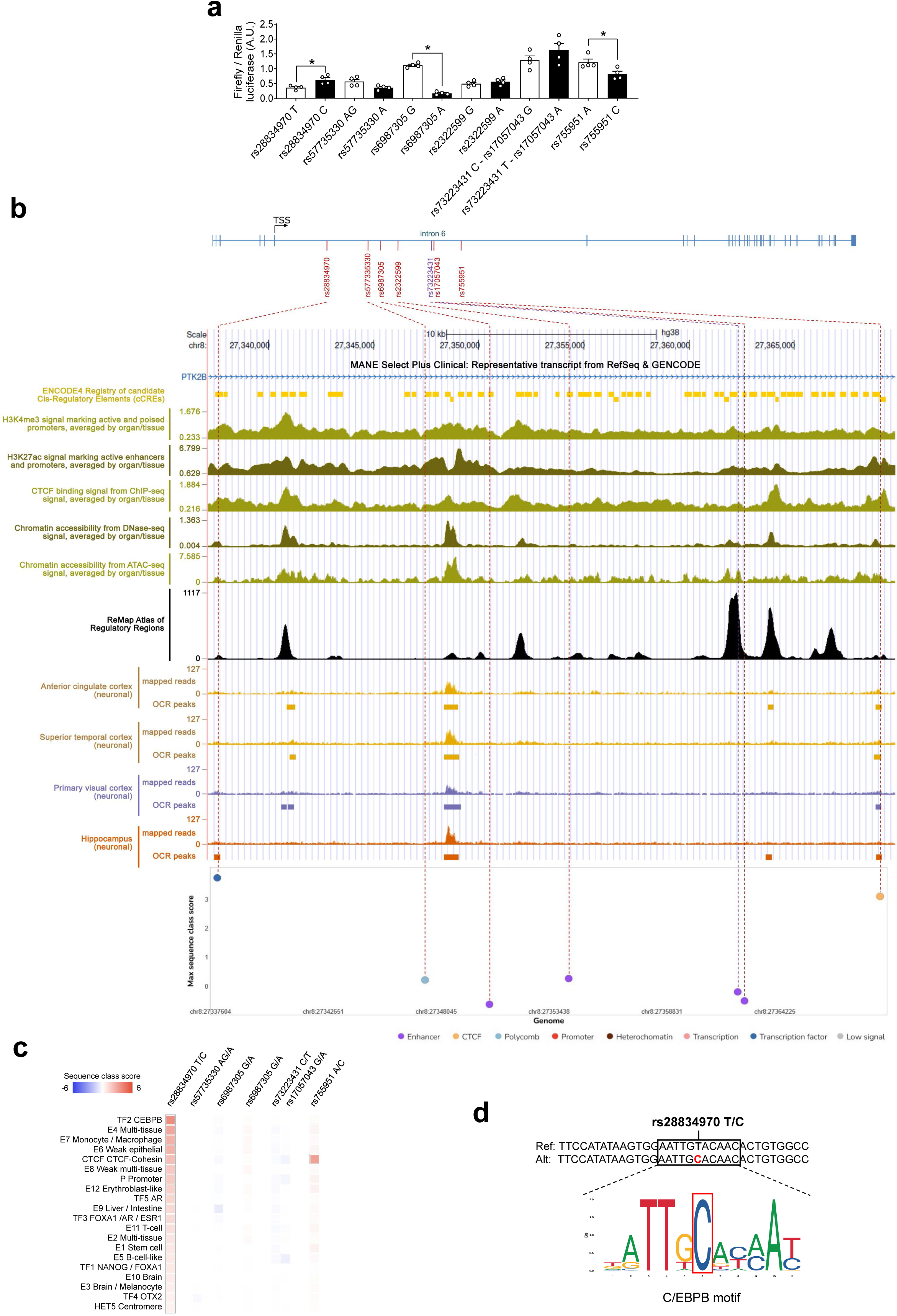
*In silico* and *in vitro* functional analysis of the seven AD-associated *PTK2B* gene variants. **(a)** Luciferase reporter assays in SH-SY5Y cells showed that three variants (i.e., rs28834970, rs6987305, and rs755951) significantly affect gene expression. Data are means ± SEM of four independent experiments performed in triplicate; Mann-Whitney U Test (two-tailed, exact p-value), * *p* < 0.05 for risk vs. common allele. **(b)** GWAS meta-analysis [2] reveals seven *PTK2B* intron 6 variants associated with AD risk. Functional genome annotation shows that rs28834970 and rs755951 are located within candidate cis-regulatory elements (cCRE) and overlap with histone marks, ATAC-seq, and DNase-seq peaks characteristic of transcriptionally active chromatin in human brain [23,24]. Both variants overlap with ChIP-seq peaks for transcriptional regulators (ReMap database) [25]. Notably, rs28834970 overlaps with ATAC-seq peaks specific to hippocampal neurons, while rs755951 overlaps with open chromatin signals in neurons from multiple brain regions [48]. The Sei deep learning algorithm predicts both variants as functional [26]. **(c)** Sei algorithm [26] predictions indicate that risk alleles for rs28834970 and rs755951 increase binding of C/EBPβ and CTCF, respectively; the remaining four variants show no predicted functional effects. **(d)** Position Weight Matrix analysis from the JASPAR 2026 database [27] indicates that the rs28834970 risk allele (C) may create a C/EBPβ transcription factor binding site within PTK2B intron 6.

To further prioritize candidates, we analyzed publicly available functional genomics datasets. ENCODE chromatin immunoprecipitation data from human brain tissues and primary cells revealed substantial enrichment of active regulatory marks (H3K4me3 and H3K27ac) and CTCF binding sites across the intronic region, consistent with active regulatory architecture (**Fig. 1b**) [23,24]. This analysis narrowed the candidate variants down to two variants, i.e., rs28834970 and rs755951, that map to candidate *cis*-regulatory elements with open chromatin signatures in DNase-seq and ATAC-seq datasets [23,24] and overlap ChIP-seq peaks from ReMap [25]. Critically, these two variants differed in brain region specificity. While rs755951 exhibited open chromatin across neurons from multiple brain regions, rs28834970 displayed open chromatin exclusively in hippocampal neurons [48]. To distinguish between them, we employed Sei [26], a machine learning framework integrating expression data, eQTL information, and evolutionary conservation to predict variant functional effects. This analysis ranked rs28834970 as the most functionally relevant variant (Sei score: 3.75 vs. 3.11 for rs755951).

The convergence of these *in vitro* and *in silico* analyses, which were in line with *PTK2B* neuronal expression, prompted us to select rs28834970 for further analysis. Supporting its potential impact in neurons, computational predictions [26,27] identified that the rs28834970 C risk allele promotes binding of the transcription factor C/EBPβ (**Fig. 1c, d**), which plays a role in memory formation and synaptic plasticity [49]. Notably, *PTK2B* has been confirmed as a direct C/EBPβ target in the mouse hippocampus [50], which is consistent with the potential hippocampus-specific regulatory profile of rs28834970.

### The rs28834970 variant impacts the regulation of *PTK2B*/PYK2 expression in hNCs

To assess the functional impact of the rs28834970 variant on the regulation of *PTK2B*/PYK2 expression in human neurons, we generated isogenic KOLF2.1J iPSCs [32] carrying homozygous rs28834970 CC genotype using CRISPR-Cas9 (**Fig. 2a**). Post-editing quality control included verification of (i) the correct incorporation of the intended mutation (**Fig. 2b**), (ii) maintenance of iPSC pluripotency, (iii) absence of both on-target and off-target editing events and (iv) absence of chromosomal aberrations (**Supplementary fig. 2**).

**Figure 2.**
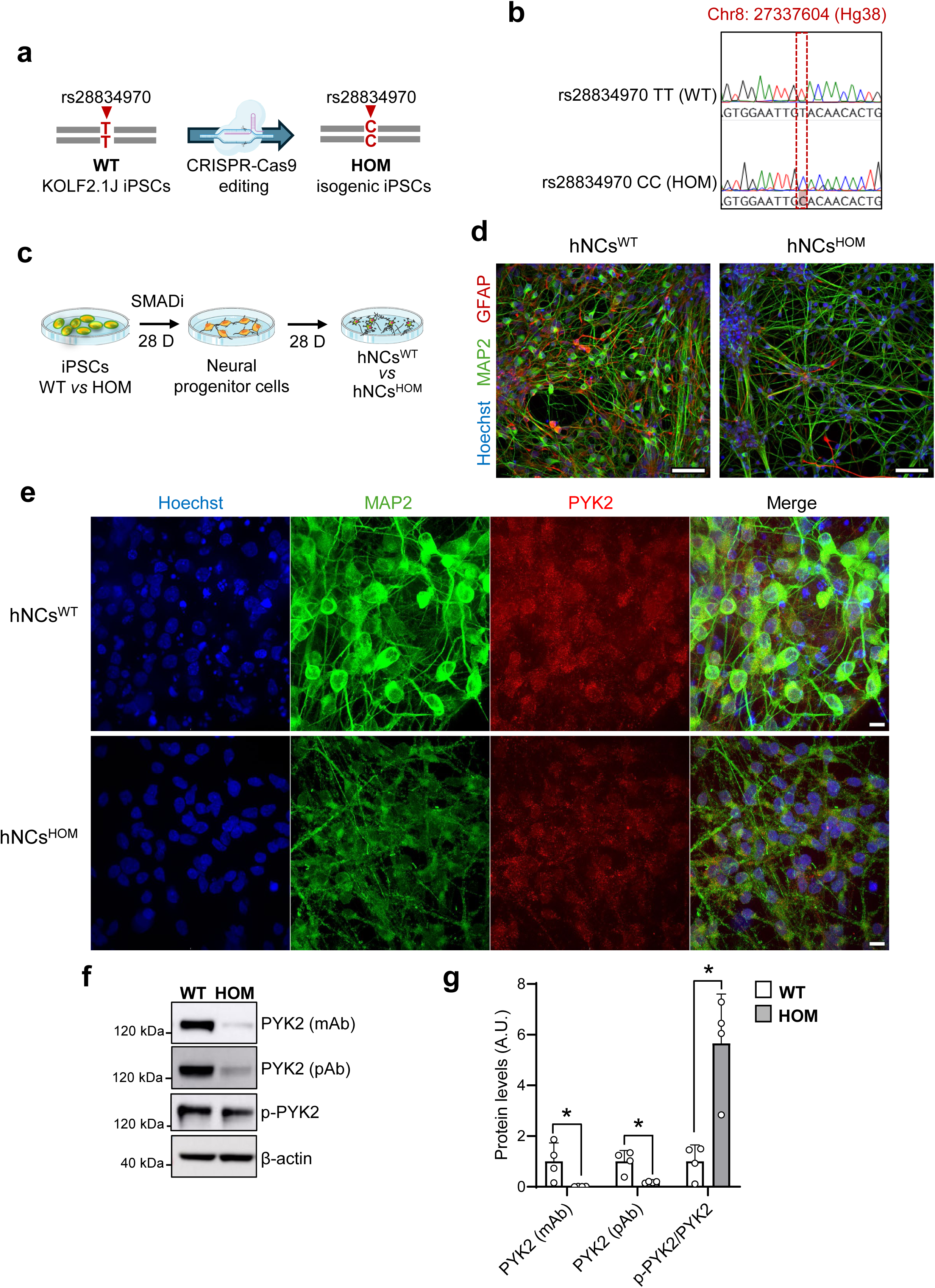
The rs28834970 variant reduces PYK2 levels in human iPSC-derived neuronal cultures. **(a)** CRISPR-Cas9 editing of KOLF2.1J iPSCs to generate isogenic lines carrying homozygous CC genotype (HOM) for rs28834970, with wild-type TT line as control (WT). **(b)** Validation of correct editing by Sanger sequencing. **(c)** Differentiation protocol for generating human neuronal cultures (hNCs) composed of forebrain neurons and astrocytes. **(d)** Immunocytochemistry confirms expression of neuronal marker MAP2 and astrocytic marker GFAP in both hNCs^WT^ and hNCs^HOM^ cultures. Scale bar = 50 µm. **(e)** Representative immunofluorescence images of 4-week-old cultures showing PYK2 immunoreactivity co-localized with neuronal marker MAP2. Scale bar = 10 µm. **(f–g)** Representative Western blots **(f)** and quantification **(g)** showing significant effects of rs28834970 on PYK2 and phospho-PYK2 Tyr402 in hNCs^HOM^ compared to hNCs^WT^. Data represent means ± SD from four independent cultures (* *p* = 0.0286, two-tailed Mann-Whitney U test).

The *in silico* and *in vitro* analyses that we conducted suggest that the impact of rs28834970 on *PTK2B* expression occurs, at least in part, in neurons. We therefore differentiated (using dual SMAD inhibition) KOLF2.1J iPSCs into human neuronal cultures (hNCs), which were composed of forebrain-like neurons and astrocytes (**Fig. 2c**). Immunocytochemical characterization of the hNCs confirmed the expression of neuronal (MAP2) and astrocytic (GFAP) markers (**Fig. 2d**). In addition, PYK2 protein was found colocalized with MAP2, which confirmed its expression within neurons (**Fig. 2e**).

First, we investigated the impact of rs28834970 on the endogenous PYK2 expression. Western blot analysis was performed using protein extracts from 4-week-differentiated hNCs carrying either the rs28834970 TT (hNCs^WT^) or the homozygous rs28834970 CC (hNCs^HOM^) risk genotype. Quantification of Western blots when using a monoclonal antibody revealed a large reduction in total PYK2 levels (‒98.9±0.7%) in hNCs^HOM^ compared to hNCs^WT^ (**Fig. 2f, g**). The strong impact of rs28834970 on PYK2 levels was further confirmed using a polyclonal antibody (‒82.2±3.3%). However, no significant change was observed in the absolute levels of phospho-PYK2 (Tyr402) in hNCs^HOM^. Nevertheless, the phospho-PYK2 to total PYK2 ratio was 5.66±1.95-fold higher than in hNCs^WT^, indicating an enrichment of the activated form of the protein (**Fig. 2f, g**).

### Transcriptomic analysis in hNCs confirmed the impact of rs28834970 on the regulation of *PTK2B* expression and suggested a role in synaptic function

We performed bulk RNA-seq in hNCs^HOM^ and hNCs^WT^ (n=5 biological replicates per condition) to evaluate the effect of rs28834970 on gene expression in our model. All samples met high-quality standards (>97% Q30 bases, >100 million mapped reads) and were aligned to the GRCh38 reference genome. DESeq2 was used for normalization and statistical testing. Principal component analysis of the top 2,000 most variable genes (**Fig. 3a**) showed clear separation between genotypes with PC1 (63% of variance) segregating samples by rs28834970 genotype and PC2 (19% of variance) showing no association with batch effects or other experimental variables. This analysis confirmed that rs28834970 genotype was the primary driver of transcriptomic variation.

**Figure 3.**
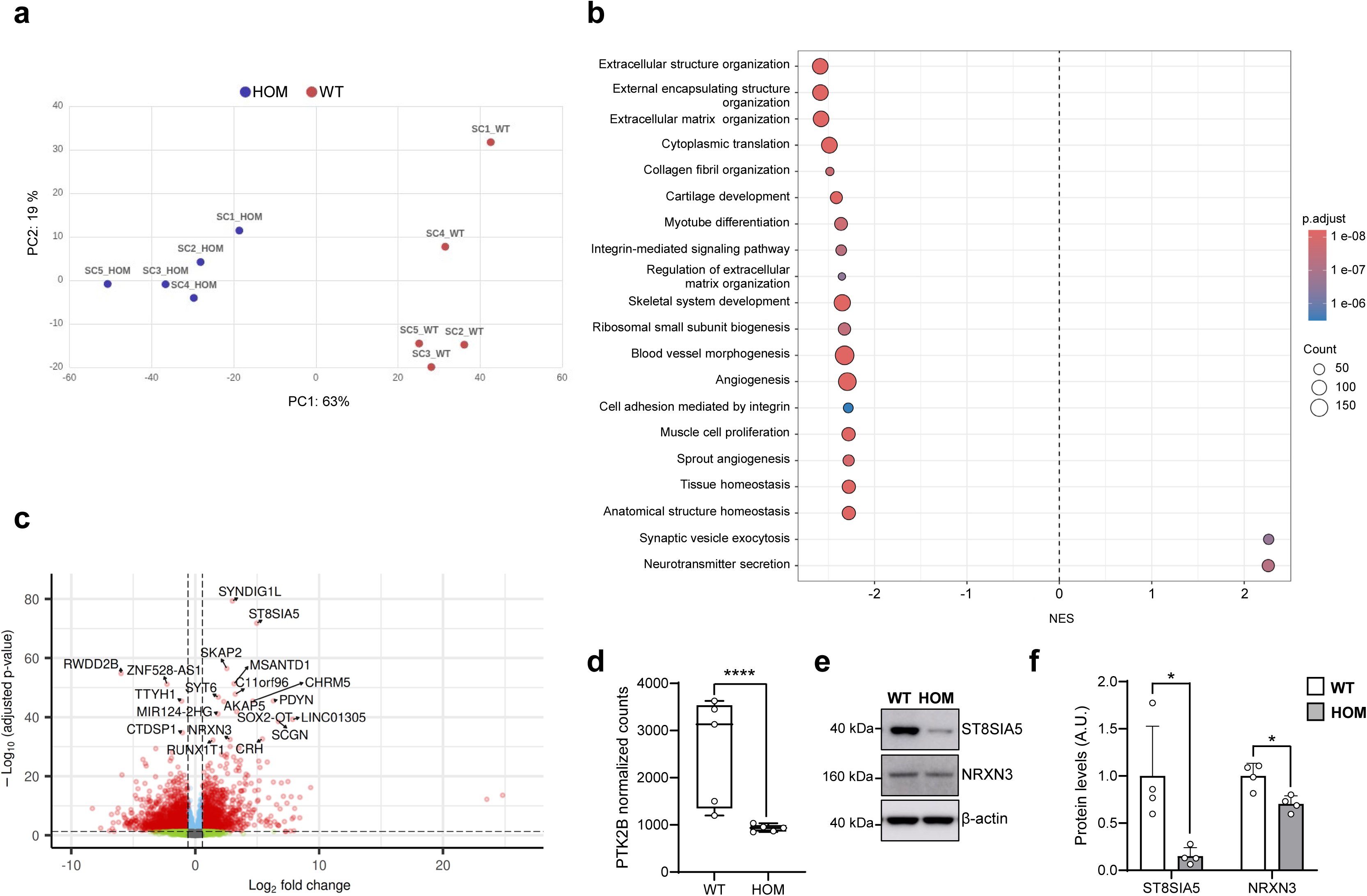
Transcriptomic analysis reveals impact of rs28834970 on synaptic function in iPSC-derived neuronal cultures. **(a)** Principal Component Analysis by genotype based on the 2,000 most variable genes; five independent cultures per genotype analyzed. The independent cultures are named SC1_HOM to SC5_HOM for hNCs^HOM^ and SC1_WT to SC5_WT for hNCs^WT^.**(b)** Gene Set Enrichment Analysis of differentially expressed genes in hNCs^HOM^ to compared to hNCs^WT^. Top 20 significantly enriched GO Biological Process gene sets (50 < size < 500, adjusted *p* < 0.05, *q* < 0.05) ranked by Normalized Enrichment Score. **(c)** Volcano plot comparing gene expression between hNCs^HOM^ and hNCs^WT^. Red dots indicate genes meeting significance thresholds (|fold change| > 1.5 and adjusted *p* < 0.05); top 20 most significant genes are labeled. **(d)** Normalized transcript counts showing significant downregulation of *PTK2B* (log₂FC = −1.5, adjusted *p* = 9.3×10^-6^) in hNCs^HOM^. **(f– g)** Representative immunoblots **(f)** and quantification **(g)** reveal reduced ST8SIA5 and NRNX3 protein levels in hNCs^HOM^ compared to hNCs^WT^. Data represent means ± SD from four independent cultures. Two-tailed Mann-Whitney U test, * *p* = 0.0286.

To identify the molecular mechanisms underlying these differences, we performed Gene Set Enrichment Analysis (GSEA) using Gene Ontology Biological Process (GO-BP) terms (**Fig. 3b and Supplementary table 2**). We identified 837 significantly enriched gene sets (p_adj_<0.05, q-value <0.05) with 724 downregulated and 113 upregulated in hNCs^HOM^ samples compared to hNCs^WT^ controls. The strongest downregulated functions were extracellular structure/matrix organization (Normalized Enrichment Score (NES)=-2.59, p_adj_=6.1×10^-9^, q=1.7×10^-9^), and cytoplasmic translation (NES=-2.49, p_adj_=6.1×10^-9^, q=1.7×10^-9^), while synaptic vesicle exocytosis (NES=2.27, p_adj_=2.5×10^-7^, q=7.0×10^-8^) and neurotransmitter secretion (NES=2.26, p_adj_=5.9×10^-8^, q=1.7×10^-8^) were significantly enhanced. These findings indicate that the rs28834970 risk genotype triggers substantial changes in extracellular matrix organization and remodeling, which is consistent with the known roles of PYK2 and its homolog FAK in integrating signals from integrins and G-protein-coupled receptors to regulate cell-matrix interactions (see [51]). The data additionally suggest that this risk genotype may impact synaptic function in human neurons.

To visualize differentially expressed genes (DEGs) between hNCs^HOM^ and hNCs^WT^, we generated a volcano plot (**Fig. 3c**). DEGs were defined with p_adj_<0.05 (**Supplementary table 3**). Of 20,205 detected transcripts, 3,944 genes met these criteria: 1,672 upregulated and 2,272 downregulated in rs28834970 HOM samples. To validate these findings, we examined mRNA levels of *PTK2B*, which showed reduced protein levels in hNCs^HOM^ (**Fig. 2f-g**). *PTK2B* mRNA was significantly decreased (log₂FC=‒1.5, p_adj_=9.3×10^-6^) (**Fig. 3d**), consistent with reduced protein levels. The two most statistically significant genes were *SYNDIG1L* (Log_2_FC=3.0, p_adj_=4.8×10^-80^) and *ST8SIA5* (Log_2_FC=5.0, p_adj_=1.6×10^-72^). *SYNDIG1L* encodes a protein that is critical for neuronal development and synaptic differentiation. *ST8SIA5* encodes α-2,8-sialyltransferase 5, which is an essential enzyme for ganglioside biosynthesis. Gangliosides have been found to be particularly abundant in neuronal membranes, and their dysregulation has been implicated in the pathology of AD [52]. *ST8SIA5* was then selected for validation at protein level *via* immunoblotting. ST8SIA5 protein levels were found to be significantly reduced in hNCs^HOM^ (‒85.0±9.1%) compared to hNCs^WT^ (**Fig. 3e, f**). We also selected *NRXN3*, the expression of which was strongly affected by the rs28834970 HOM genotype (Log2FC=2.8, adjusted p=3.3×10^-33^). NRXN3 is a cell adhesion molecule that controls presynaptic and postsynaptic properties of synaptic transmission, and has been associated with AD [53–55]. While *NRXN3* transcripts were up-regulated, we observed decreased NRXN3 protein levels (‒29.7±8.6%) (**Fig. 3e, f**).

### rs28834970 alters neuronal bursting and impacts synaptic structure in hNCs

Our transcriptomic analyses indicate an impact of *PTK2B* underexpression on synapse-related pathways. To assess the impact of the rs28834970 risk genotype on synaptic function in hNCs, we performed longitudinal multielectrode array (MEA) recordings in 48-well plates during neuronal differentiation for four weeks (**Fig. 4**). Compared to hNCs^WT^ cultures, hNCs^HOM^ cultures exhibited progressive alterations in network activity, with the most pronounced differences observed at days 21 and 28 of differentiation (**Fig. 4a**). The number of active electrodes was reduced in hNCs^HOM^ compared to hNCs^WT^ from day 14 (2.8±1.9 *vs*. 7.5±4.1, respectively, p<0.0001) through day 28 (1.7±1.5 *vs.* 4.8±4.1, respectively, p<0.0001) (**Fig. 4b**). In contrast, mean firing rate (**Fig. 4c**) and the number of bursts per electrode (**Fig. 4d**) did not differ significantly between genotypes across the differentiation time course. However, analysis of burst characteristics revealed genotype-dependent alterations in neuronal bursting. At day 21, hNCs^HOM^ cultures exhibited an increased burst frequency (0.17±0.13 Hz in hNCs^HOM^ *vs*. 0.12±0.08 Hz in hNCs^WT^, adjusted p=0.0271, **Fig. 4e**) accompanied by a reduced number of spikes per burst (8.6±3.4 in hNCs^HOM^ *vs.* 10.8±5.1 in hNCs^WT^, adjusted p=0.0339, **Fig. 4f**). At day 28, burst duration was increased in hNCs^HOM^ compared to hNCs^WT^ cultures (0.32±0.12 s *vs.* 0.25±0.06 s, p=0.0366, **Fig. 4g**), together with prolonged interspike interval within bursts (0.05±0.01 s *vs.* 0.04±0.01 s, respectively, p=0.0097, **Fig. 4h**), while the interburst interval did not change (**Fig. 4i**). Thus, although overall firing rates and burst occurrence were largely preserved, the rs28834970 risk genotype was associated with reduced network recruitment and altered temporal organization of neuronal bursting during network maturation.

**Figure 4.**
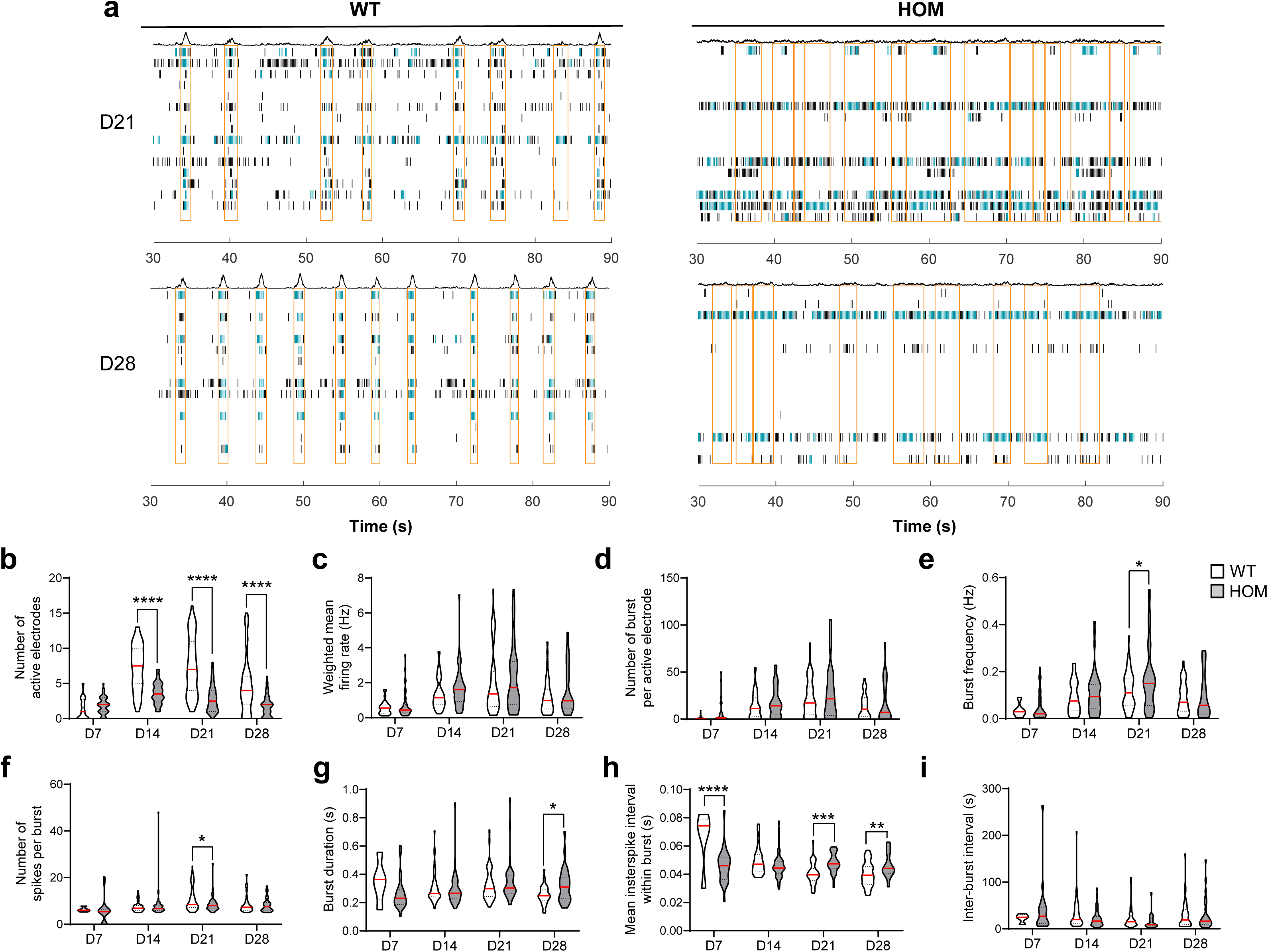
Longitudinal microelectrode array recordings show altered network activity in hNCs carrying the risk genotype for rs28834970. **(a)** Representative raster plots of action potentials recorded from hNCs^WT^ and hNCs^HOM^ cultures at DIV21 and DIV28. Each row represents a single electrode, and each black bar indicates the start of an action potential. Spikes in bursts are indicated as green bars and network bursts are indicated by orange rectangles. **(b)** Violin plots quantifying the number of active electrodes per condition across development (DIV7, DIV14, DIV21, and DIV28). **(c–i)** Quantitative analysis of network electrophysiological properties including weighted mean firing rate **(c)**, number of bursts per active electrode **(d)**, burst frequency **(e)**, number of spikes per burst **(f)**, interburst interval **(g)**, burst duration **(h)**, and interspike interval within bursts **(i)** across all developmental timepoints. Recordings were analyzed using mixed-effects models with Holm-Šídák’s multiple comparisons test to account for repeated measures across development. N = 56 wells in hNCs^WT^ *vs* n = 56 wells in hNCs^HOM^ from 3 independent cell cultures.* *p* < 0.05, ** *p* < 0.01, *** *p* < 0.001, **** *p* < 0.0001.

To assess whether rs28834970 affects synaptic loss, we quantified synaptic density using immunocytochemistry with antibodies targeting the presynaptic marker Synaptophysin and the postsynaptic marker Homer1 in microfluidic devices (**Fig. 5a-b**). Synapse density was calculated by quantifying Homer1 puncta in spatial proximity of Synaptophysin puncta using custom Matlab analysis [42]. This analysis revealed that hNCs^HOM^ exhibited a significantly lower dendritic network volume compared to hNCs^WT^ as revealed by MAP2 staining (33.3% decrease, p=0.0012, **Fig. 5c**), which was accompanied by a significant decrease in the number of Homer1 puncta (27.0%, p=0.0121, **Fig. 5d**), but not in the number of Synaptophysin puncta (8.7% decrease, p=0.166, **Fig. 5e**). This differential impact resulted in a significantly lower fraction of Synaptophysin puncta assigned by at least one Homer1 puncta in hNCs^HOM^ compared to hNCs^WT^ (11.7% decrease, p=0.0220, **Fig. 5f**) and translated to a significant decrease in the number of synapses, as defined as the number of Synaptophysin puncta with Homer1 assignments (18.9% decrease, p=0.0303, **Fig. 5g**). Normalizing these read-outs by the MAP2 volume, one can obtain relative changes in the densities of pre- and postsynaptic puncta and of synapses. Compared to hNCs^WT^, hNCs^HOM^ exhibited a relatively higher Synaptophysin spot density (1.35-fold increase, p=0.0220, **Fig. 5h**), but the relative densities of Homer1 puncta and of synapses remained unchanged (p=0.8408 and p=0.1541, respectively, **Fig. 5i-j**). Overall, these results indicate that rs28834970 has a detrimental impact on dendritic/postsynaptic maturation, decreasing synaptic connectivity and number of synapses, but not necessarily their relative density, which is consistent with the decreased number of active MEA electrodes and with the altered bursting behavior.

**Figure 5.**
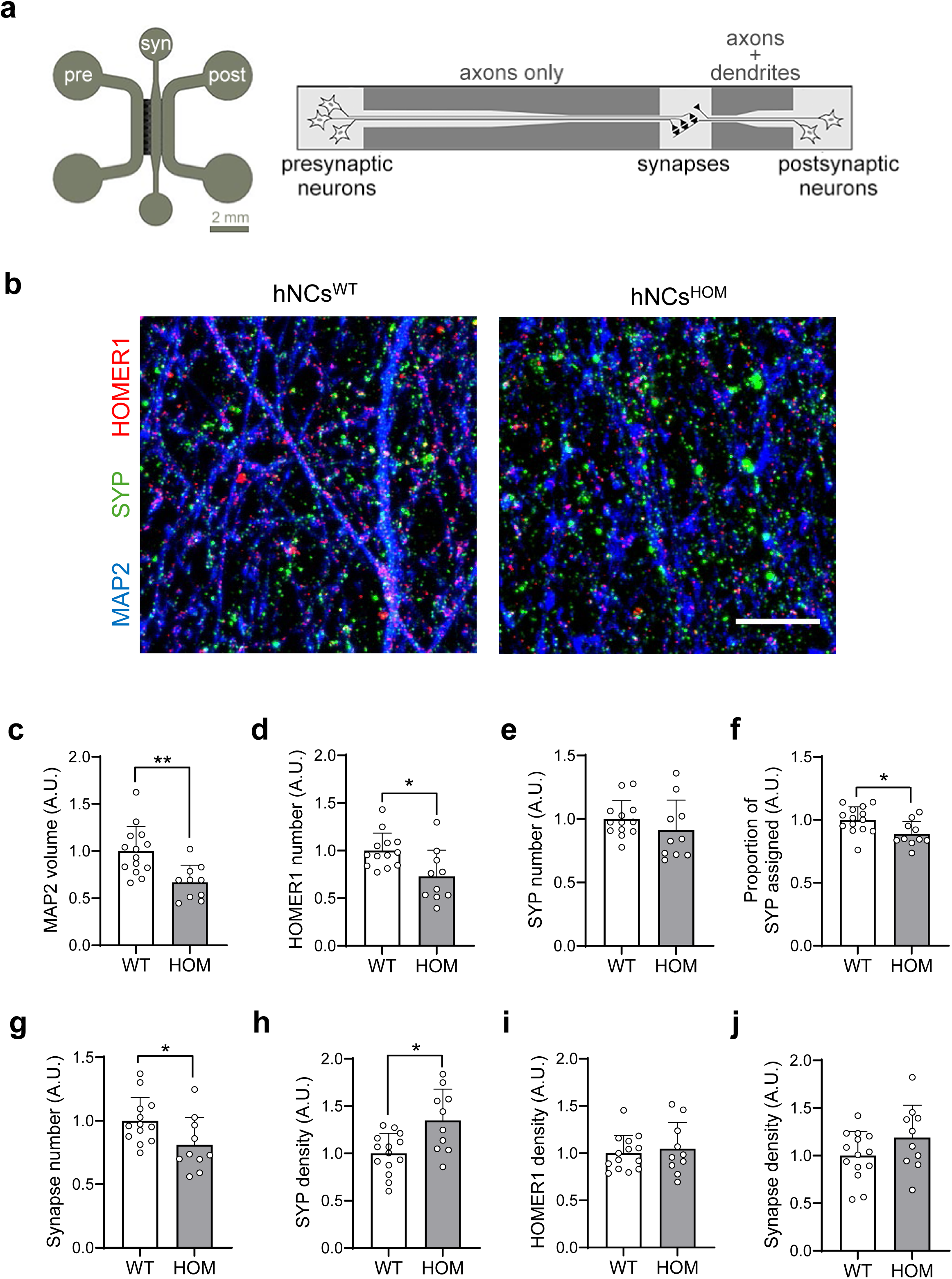
Analysis of synaptic structure in microfluidic devices. **(a)** Schematic depiction of the trichamber microfluidic device that isolates neurites from their cell bodies. The graphic on the right is not drawn to scale. **(b)** Representative immunofluorescence images of the synaptic chamber showing Synaptophysin (SYP, green) and Homer1 (red) puncta and dendritic network by MAP2 (blue). Images were acquired as *z*-stacks of 0.2 µm, deconvoluted, and subjected to maximum intensity projection. **(c-j)** Quantitative analysis of synaptic connectivity showing MAP2 volume **(c)**, SYP spot number **(d)**, Homer1 spot number **(e)**, fraction of SYP spots assigned by at least one Homer1 spot **(f)**, synapse number (number of SYP spots with Homer1 assignments, **g**), SYP and Homer1 spot densities (spot numbers per MAP2 volume, **h-i**), and synapse density (synapse number per MAP2 volume, **j**). N = 3 independent experiments each with 2−6 devices per condition. Data were normalized by the mean of the WT group in each experiment before pooling the experiments. Two-tailed Mann-Whitney U test, * *p* < 0.05, ** *p* < 0.01, *** *p* < 0.005.

### The rs28834970 risk genotype increases Tau phosphorylation and APP processing, potentially through GSK3β activation

Beyond its involvement in synaptic structure and function, we tested whether the rs28834970 risk allele could recapitulate AD hallmarks in hNCs. Western blot quantification revealed no significant changes in total Tau levels in hNCs^HOM^ compared to hNCs^WT^ (**Fig. 6a, b**). In contrast, Tau phosphorylation was significantly elevated at AD-associated sites Thr181 (1.84±0.52-fold), Thr217 (2.45±0.23-fold), Thr231 (1.70±0.46-fold) and Ser396-Ser404 (1.81±0.50-fold) in hNCs^HOM^ (**Fig. 6a, b**). We observed also a massive increase in Tau phosphorylation at Ser202/Thr205 that resulted in a binary ‘all-or-nothing’ signal, making standard densitometry impossible (**Fig. 6a**). To investigate whether this effect involved key Tau kinases, we assessed total and phosphorylated levels of ERK1/2, p38-MAPK, AKT, and GSK3β *via* immunoblotting (**Fig. 6c-f**). The risk genotype for rs28834970 did not significantly impact total AKT or GSK3β protein levels. However, it significantly decreased total ERK1/2 and p38 MAPK levels by 37.1 ± 21.0% and 49.5 ± 5.5%, respectively, while also reducing absolute levels of phosphorylated ERK2 (Thr202/Tyr204) by 38.7 ± 11.4%. Phosphorylated ERK1 exhibited a comparable, albeit non-significant, decrease of 43.4 ± 21.0% (p=0.0571). The ratios of the phosphorylated-to-total forms of ERK1/2, AKT and p38 MAPK remained unchanged. In contrast, GSK3β phosphorylation at Tyr216 increased 1.34-fold in hNCs^HOM^ compared to hNCs^WT^ (p=0.0286).

**Figure 6.**
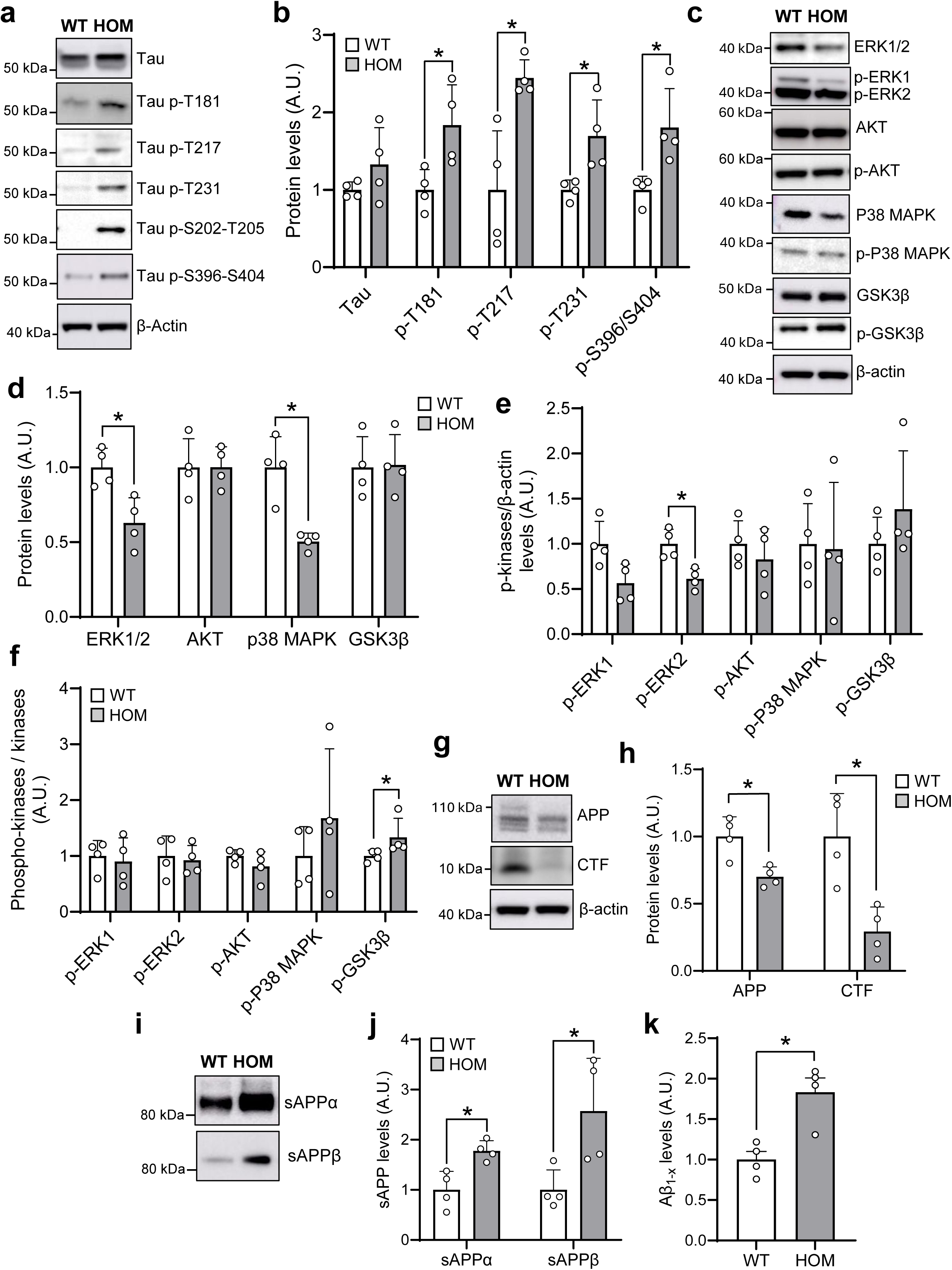
The rs28834970 risk genotype enhances Tau phosphorylation and APP processing in iPSC-derived neuronal cultures. Representative western blots **(a)** and quantification **(b)** showing increased phorphorylation of Tau at Thr181, Thr217, Thr231, Ser202/Thr204, and Ser396/Ser404 in hNCs^HOM^ compared to hNCs^WT^ (normalized to total Tau). **(c-f)** Changes in Tau kinases (p38-MAPK, ERK1/2, AKT and GSK3β) and their phosphorylated (activated) forms in hNCs^WT^ *vs* hNCs^HOM^. **(g,h)** Representative western blots **(g)** and quantification **(h)** showing decreased total APP, and CTF levels in hNCs^WT^ *vs* hNCs^HOM^. **(i-j)** Representative western blots **(i)** and quantification **(j)** showing increased sAPPα and sAPPβ levels in conditioned media of hNCs^HOM^. **(k)** Alpha-LISA quantification of elevated Aβ peptide levels in hNCs^HOM^ conditioned media. sAPPα, sAPPβ and Aβ peptide measurements were normalized to total protein levels. Data represent means ± SD from four independent cell cultures Two-tailed Mann-Whitney U test, * *p* = 0.0286.

Western blot analysis of APP revealed a significant 30.0±15.6% decrease in total APP in hNCs^HOM^ compared to hNCs^WT^ (**Fig. 6g, h**), suggesting either impaired production or accelerated metabolism. To distinguish between these possibilities, we quantified APP C-terminal fragments (CTFs) in cell lysates and secreted APP metabolites (sAPPα, sAPPβ, and Aβ_1-x_) in the conditioned media. CTFs were significantly reduced (−76.7±8.3%) in hNCs^HOM^ (**Fig. 6g, h**). In contrast, the levels of both sAPPα and sAPPβ (as measured by Western blotting, **Fig. 6i, j**) and of Aβ_1-x_ (measured by AlphaLISA, **Fig. 6k**) were significantly higher in hNCs^HOM^ than in hNCs^WT^ (1.78±0.20-fold, 2.57±1.05-fold and 1.83±0.18-fold, respectively), indicating enhanced APP processing through both non-amyloidogenic and amyloidogenic pathways.

To independently validate these findings, we leveraged previously published data from a genome-wide, high-content siRNA screening approach designed to identify regulators of APP metabolism [56]. This screen employed HEK293 cells stably expressing an mCherry-APP695-YFP fusion construct, which enables rapid detection and quantification of APP processing events through fluorescence intensity changes (**Supplementary fig. 3**). In this screen, *PTK2B* silencing resulted in significantly increased levels of mCherry (1.62±0.46-fold) and YFP (2.05±1.05-fold) fluorescence, indicating that PYK2 downregulation enhances APP metabolism in this model. This is consistent with the phenotypes observed in hNCs. Collectively, these findings demonstrate that the rs28834970 CC risk genotype elevates Tau pathological phosphorylation while simultaneously accelerating APP metabolism and promoting Aβ production, thereby mechanistically linking this genetic variant to increased AD susceptibility.

## DISCUSSION

In a human iPSC-derived neuronal model, we demonstrated that the rs28834970 risk genotype substantially reduces PTK2B/PYK2 levels. Transcriptomic analyses indicate that rs28834970 impacts pathways related to extracellular matrix organization and synaptic function. The latter finding was supported by immunocytochemistry and MEA recordings demonstrating that rs28834970 alters synaptic organization and neuronal electrical activity. Finally, we demonstrated that this AD risk variant also impairs both Tau phosphorylation and APP processing. Collectively, our findings establish a link between *PTK2B*/PYK2 transcriptional dysregulation driven by genetic variation and AD-related pathogenic processes.

Interestingly, the C risk allele of rs28834970 has previously been shown to bind preferentially C/EBPβ and decrease *PTK2B* mRNA levels in iPSC-derived macrophages and microglia [57]. However, no impact of this allele was detected at the PYK2 protein level in these cell types. In contrast, our results consistently showed that the homozygous risk genotype, which is present in 13.7% of European populations, reduced both *PTK2B* mRNA and PYK2 protein levels by over 80% in human neuronal cultures. When overexpressed and activated, C/EBPβ is known to drive neuroinflammation, Aβ production, Tau pathology, and disease progression [58,59]. In an AD mouse model, intracellular truncated APP C586-695 induced nuclear translocation and target binding of C/EBPβ [60], whereas *PTK2B* was found to be overexpressed in the hippocampus of C/EBPβ-knockout mice [50]. Together, these observations support the hypothesis that *PTK2B*/PYK2 plays cell-type-specific roles across neurons, astrocytes, and microglia, framing the rs28834970-impacted C/EBPβ regulatory axis as a central node in AD pathogenesis.

Transcriptomic analysis pointed to dysregulation in synaptic transmission pathways caused by the AD risk variant rs28834970. Functional synaptic impairment was corroborated by our MEA findings in hNCs. The preservation of mean firing rates alongside disrupted burst architecture in rs28834970 risk-homozygous cultures indicates that while neurons remain electrophysiologically active, their network coordination is compromised during maturation. This electrophysiological phenotype likely reflects multiple converging mechanisms: (j) reduced synaptic connectivity or strength, accounting for persistent decreases in active electrode counts and spikes per burst, (ii) impaired temporal coordination of firing, evidenced by elevated burst frequency and prolonged interspike intervals; and (iii) altered network rhythmicity, leading to prolonged bursts. These network alterations suggest that rs28834970 impairs functional maturation and network assembly rather than inducing non-specific suppression of intrinsic neuronal excitability. Immunocytochemistry in microfluidic devices revealed that rs28834970 impacts dendritic development and postsynaptic maturation, consistent with these findings. Given the established functions of PYK2 in neurite development, NMDAR-dependent signaling, and activity-dependent synaptic plasticity [11], attenuated *PTK2B*/PYK2 activity provides a plausible mechanistic bridge between the rs28834970 variant and circuit-level dysfunction. Neuronal network desynchrony is a recognized feature of early AD progression [61]. Disrupted burst dynamics and network recruitment in risk-genotype cultures may therefore represent an early functional vulnerability that lowers the threshold for subsequent pathological processes. Interestingly, *NRXN3* was among the most significantly upregulated mRNA in our transcriptomic analysis, yet NRXN3 protein levels decreased in rs28834970-homozygous hNCs. Remarkably, this inverse mRNA/protein signature mirrors our previous observations for the AD risk factor *PLCG2* in hNCs, where genetically driven *PLCG2* downregulation caused synaptic dysfunction, GSK3β activation, Tau hyperphosphorylation, elevated Aβ production, and decreased NRXN3 protein despite elevated *NRXN3* mRNA levels [34].

Although total PYK2 levels were markedly reduced in the presence of the rs28834970 risk genotype, the relative proportion of its phosphorylated form (Tyr402) was increased, suggesting a potential compensatory mechanism that enhances residual PYK2 activity. In hNCs, the rs28834970 risk genotype increased Tau phosphorylation across multiple pathogenic epitopes, including Thr181, Thr217, Thr231, Ser202/Thr204, and Ser396/Ser404. These findings align with previous *in vitro* data demonstrating that both *PTK2B*/PYK2 overexpression and inhibition enhance Tau phosphorylation at Ser396/Ser404 and Ser202/Thr204 [20]. To identify the kinases responsible for increased Tau phosphorylation, we examined major Tau kinases. The rs28834970 risk genotype significantly reduced total ERK1/2 and p38 MAPK protein levels without altering the phosphorylation ratio (p-kinases/total kinases) or mRNA expression (as observed in our RNA-seq data), indicating a post-transcriptional mechanism. Accordingly, the selective reduction in total MAPK (i.e. ERK1/2 and p38 MAPK) protein abundance, coupled with preserved phosphorylation ratios, suggests potential mechanisms of: (i) reduced protein synthesis as suggested by our transcriptomic analysis; (ii) enhanced degradation; and (iii) altered stability, rather than impaired MAPK activation. This finding argues against MAPK as the primary driver of increased Tau phosphorylation in hNCs carrying the rs28834970 risk genotype. A more direct mechanism emerged from our finding that rs28834970 increased GSK3β phosphorylation at Tyr216, a residue within the catalytic domain that enhances kinase activity, positioning GSK3β as a potential mediator of the Tau hyperphosphorylation phenotype. The functional relationship between PYK2 and GSK3β in the central nervous system is well documented and involves a direct kinase– substrate interaction linked to calcium signaling and Tau phosphorylation [62,63]. Neuronal PYK2 phosphorylates and activates GSK3β at Tyr216 downstream of G-protein–coupled receptor activation: lysophosphatidic acid stimulates phospholipase C-mediated PIP2 hydrolysis, triggering intracellular calcium release that activates PYK2 [63]. Activated PYK2 physically associates with GSK3β in stable protein complexes to phosphorylate it at Tyr216. Once activated, GSK3β hyperphosphorylates Tau at key pathogenic epitopes (including Ser202/Thr204 and Ser396/Ser404) [64]. Additionally, our results demonstrate that rs28834970 alters APP processing in hNCs, resulting in lower levels of full-length APP and CTF, and higher levels of sAPPα, sAPPβ and Aβ. GSK3β is known to modulate APP metabolism and promote Aβ generation, though its role in amyloidogenesis is less known than in Tau phosphorylation (reviewed in [65]). Transgenic mice conditionally overexpressing GSK3β in the forebrain display increased APP carboxy-terminal fragment cleavage, elevated Aβ production, and accelerated amyloid plaque deposition, whereas GSK3β inhibition mitigates plaque burden and cognitive deficits [66]. Elevated GSK3β activity also disrupts neuronal architecture and function; GSK3β-overexpressing mice display abnormal neuronal maturation and dendritic atrophy [67], while rats expressing constitutively active GSK3β exhibit impaired oscillatory dynamics in the prefrontal cortex and hippocampus [68]. Finally, sporadic AD iPSC-derived neurons recapitulate this triad of Tau hyperphosphorylation, elevated Aβ levels, and sustained GSK3β activation [69], matching the cellular phenotype observed in our rs28834970 risk hNCs.

In previous *in vivo* studies, PYK2 has displayed a complex, context-dependent role in AD pathophysiology, with seemingly disparate findings regarding neuronal deficits, amyloid toxicity, and Tau pathology. On one hand, PYK2 appears to mediate Aβ oligomer-induced synaptotoxicity. Salazar *et al*. [13] showed that crossing APP/PS1 mice with *Pyk2*-KO mice rescued cognitive deficits. Similarly, *Pyk2* deletion prevented Aβ-induced hippocampal neuronal hyperactivity and spine loss, whereas neuronal Pyk2 overexpression reduced dendritic spine density via its proline-rich domain independently of catalytic activity [14]. Conversely, crossing 5XFAD mice with *Pyk2*-deficient mice failed to improve their phenotype [17]; instead, 5XFAD mice exhibited a functional deficit in Pyk2 that was rescued by AAV-mediated Pyk2 overexpression. Consistent with a neuroprotective role for functional PYK2, genetic reduction [70] or pharmacological inhibition [71] of STEP (the primary tyrosine phosphatase that inactivates PYK2 in neurons) restored cognitive function in 3xTg-AD mice, and postsynaptic PYK2 overexpression protected primary hippocampal neurons against Aβ-induced synaptotoxicity [42]. Regarding Tau pathology, the *Drosophila* ortholog of PYK2 interacts with Tau to modulate its toxicity [18], and Pyk2 co-localizes with Tau at murine hippocampal synapses, where *Pyk2* knockout reduces Tau synaptic localization [14]. Furthermore, *Pyk2* deletion in a mouse tauopathy model accelerated Tau phosphorylation, decreased survival, and worsened memory deficits [20]. In human AD brains, hyperphosphorylated Tau co-localizes with PYK2 [18], and total PYK2 levels were reduced whereas phosphorylated PYK2 at Tyr402 was enhanced compared to controls [42]. Collectively, these findings suggest that both severe deficit and excessive overexpression of *PTK2B*/PYK2 disrupt homeostatic signaling. Our findings in human isogenic hNCs carrying the rs28834970 variant indicate that *PTK2B* downregulation does not merely alter neuronal sensitivity to pre-existing pathology, but can directly initiate AD-like amyloid, Tau, and network abnormalities. Several limitations of this study should be noted. First, 2D iPSC-derived neurons display variable maturation depending on the differentiation protocol, lack the spatial organization of mature neural circuits, and lack systemic organ-level interactions that influence AD *in vivo*. These findings should be validated in 3D brain organoids, assembloids, or humanized knock-in animal models. Second, our study did not evaluate whether rs28834970 alters early neurodevelopmental trajectories or if observed phenotypes reflect developmental vulnerability rather than adult-onset neurodegeneration. Third, because PYK2 is expressed in astrocytes and astrocytes regulate synaptic maintenance, Aβ clearance, and Tau pathology, the precise cell-autonomous versus non-cell-autonomous contributions of astrocytes to the rs28834970 phenotype remain to be determined.

## CONCLUSION

In conclusion, the dysregulation of *PTK2B* caused by the rs28834970 variant may drive multiple pathological processes involved in AD, including alterations in synaptic structure and function, Tau hyperphosphorylation, and increased APP processing. These findings validate the use of human iPSC-derived neuronal cultures as a preclinical model for the functional assessment of AD-associated variants and targeted therapeutic screening against the *PTK2B*/PYK2 signaling axis. Future studies should: (i) extend these findings to humanized rs28834970 knock-in mouse models; (ii) assess cell-type-specific contributions using isolated neuronal, microglial and astrocytic models; and (iii) identify specific downstream effectors of PYK2 that could be targeted therapeutically in individuals carrying the rs28834970 variant.

## Availability of data and materials

The RNA-seq data will be uploaded to a public repository prior to publication.

## Authors’ contributions

Conceptualization: NLW, JD; Methodology: NLW, MH, FL, BGB, DK, DP, JCL, JD; Formal analysis and investigation: NLW, MH, FL, ML, VG, FM, KG, CH, AC, VB, FR, NM, PW, SDS, LMM, AMF, CN, AD, JGG, MM, JC; Writing - original draft preparation: NLW, JCL, JD; Writing - review and editing: BGB, JC, DK, DP; Funding acquisition: MRC, PA, DP, DK, JCL, JD; Supervision: DP, DK, JCL, JD. All authors read and approved the manuscript.

## Acknowledgments

We thank the high-performance computing service at the University of Lille. We thank the BioImaging Center Lille (BICeL), part of Plateformes Lilloises en Biologie et Santé (PLBS) - UAR 2014 - US 41, for microscopy resources. We thank Karine Blary at the IEMN Lille for the microfabrication work.

