## Supplementary figures for "The Alzheimer’s disease-associated rs28834970 variant in *PTK2B* impairs electrical activity and regulates Tau phosphorylation and APP processing in human neuronal cultures"

### SUPPLEMENTARY FIGURE LEGENDS

#### Supplementary Figure 1. *PTK2B* expression in single-nucleus RNA-seq datasets from Alzheimer's disease brains

**(a–c)** *PTK2B* expression percentile among seven cell types ranked by mRNA abundance in the entorhinal cortex **(a)**, superior frontal gyrus **(b)**, and temporal cortex **(c)**. Micro, microglia; Exc, excitatory neurons; Inh, inhibitory neurons; Astro, astrocytes; Oligo, oligodendrocytes; OPC, oligodendrocyte precursor cells; Endo, endothelial cells; Epi, epithelial cells. **(d)** Forest plot summarizing donor-level associations between *PTK2B* expression and Continuous Pseudoprogression Score (CPS) in the SEA-AD middle temporal gyrus dataset across cell types. Regression coefficients ($\beta$) and 95% confidence intervals are shown; positive $\beta$ indicates increased *PTK2B* expression with elevated CPS. Point size reflects number of donors; color represents statistical significance after correction for multiple testing ($-\log_{10}$ FDR; Benjamini-Hochberg procedure). **(e–g)** Scatter plots showing donor-level mean *PTK2B* expression versus CPS with linear regression fit (blue line with 95% confidence interval shaded) and LOESS (Locally Estimated Scatterplot Smoothing) curve (red line) for visual assessment of model fit. Linear regression results by cell type are reported as $R^2$ values and FDR-adjusted p-values, with significance determined at $p < 0.05$.

#### Supplementary Figure 2. Quality control of CRISPR-edited KOLF2.1J iPSCs for the *PTK2B* rs28834970 variant

**(a)** Pluripotency confirmed by immunofluorescence of SOX2, OCT4, and SSEA4 markers in both WT (rs28834970 TT) and HOM (rs28834970 CC) edited iPSCs. Scale bars = 100 μm. **(b)** PCR fingerprinting at the human D1S80 locus (chromosome 1) confirms genetic identity between WT and HOM iPSCs relative to controls (CTL1, human genomic DNA #1; CTL2, human genomic DNA #2; CTL3, HEK293 genomic DNA). **(c–d)** Absence of unintended on-target effects verified by qPCR **(c)** and heterozygous SNP genotyping of the edited *PLCG2* locus by whole-genome sequencing **(d)**. ID, variant identifier (chr:position:reference\_allele:alternate\_allele, GRCh38 assembly); CHROM, chromosome; POS, position; REF, reference allele; ALT, alternate allele; GT, genotype (0, reference; 1, alternate); AD, allelic depth; GQ, Phred-scaled genotype quality (maximum 99). **(e–f)** qPCR **(e)** and WGS **(f)** analyses confirmed the absence of CRISPR-induced chromosomal aneuploidy or structural variants.

#### Supplementary Figure 3. High content siRNA screening reveals impact of *PTK2B* downregulation on APP processing

**(a)** Schematic representation of APP with inserted fluorophores (mCherry and YFP) used in the high-content siRNA screening. **(b)** Representative fluorescence microscopy images and quantification demonstrating the effect of *PTK2B* silencing on mCherry and YFP intensity. Scale bar = 20 μm. **(c)** Graphs showing variations in mean fluorescence intensity per cell ($n > 300$) following transfection of HEK293-APP-mcherry-YFP cells with siPTK2B versus non-targeting siRNA control (siNT) across three independent experiments.

Supplementary Figure 1

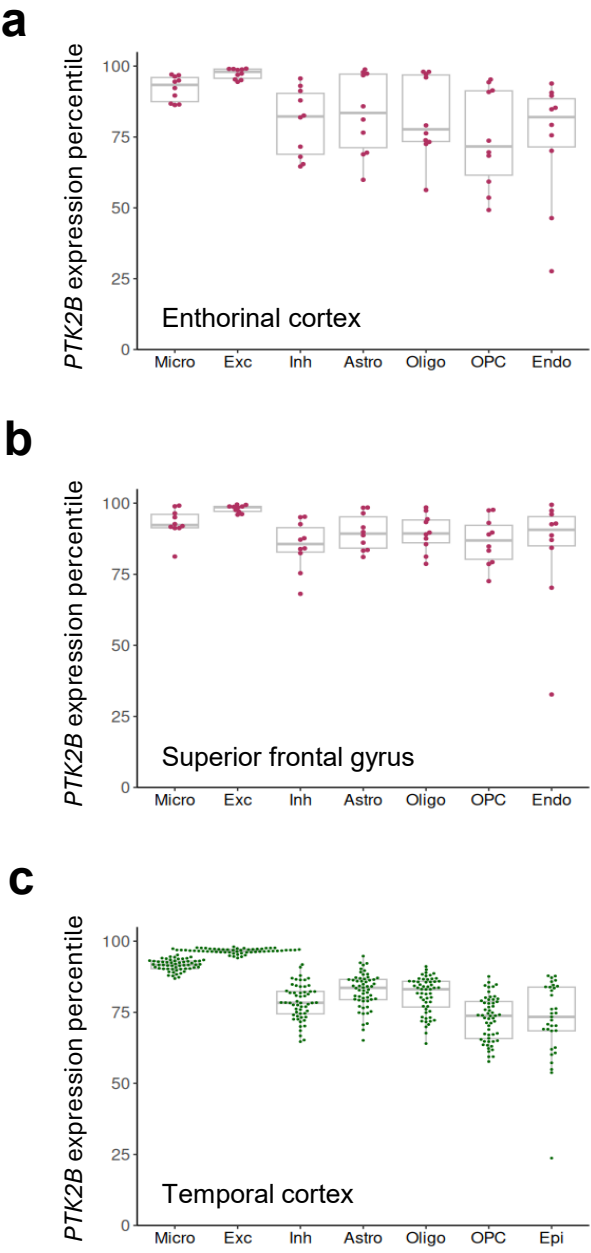

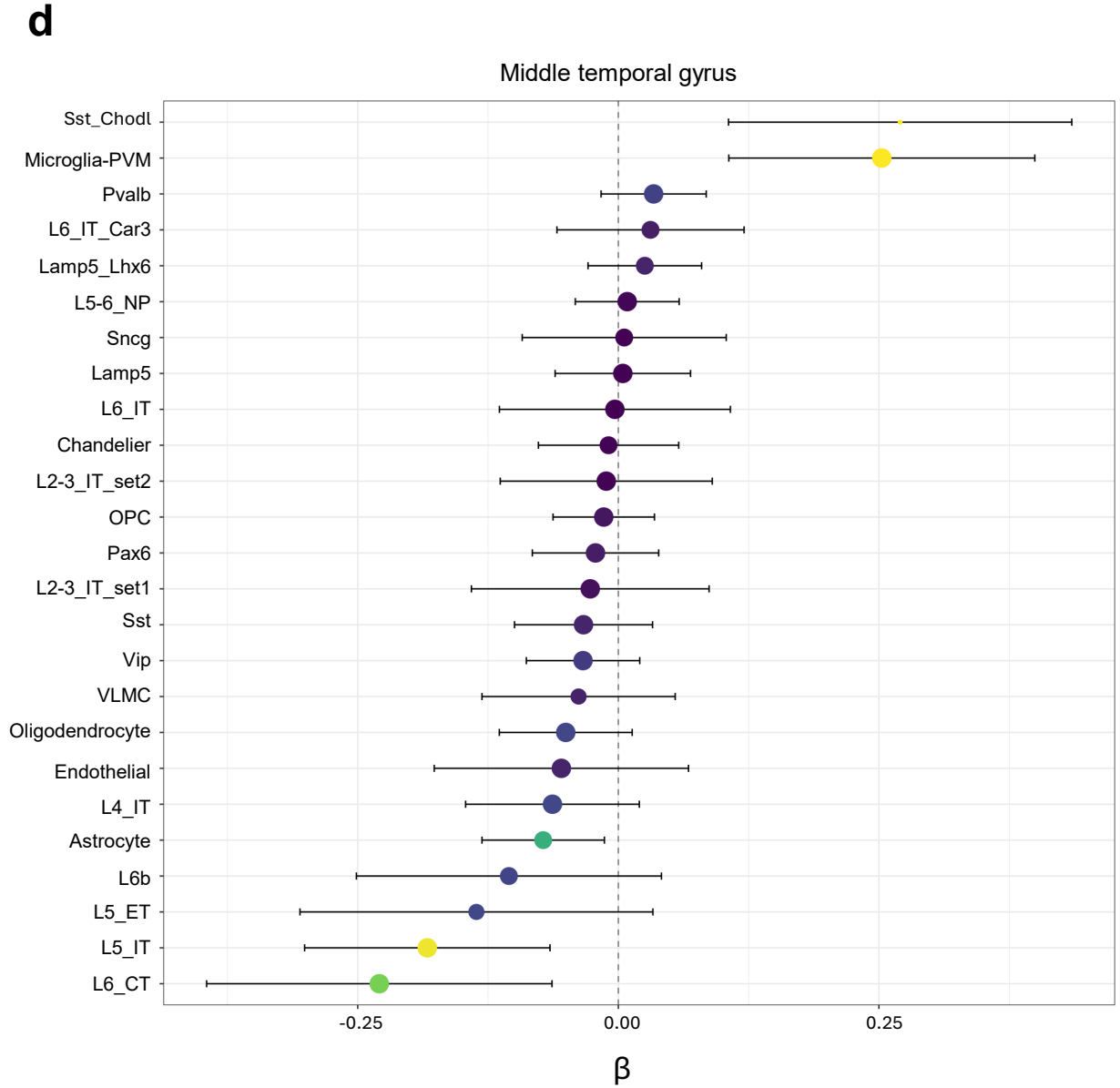

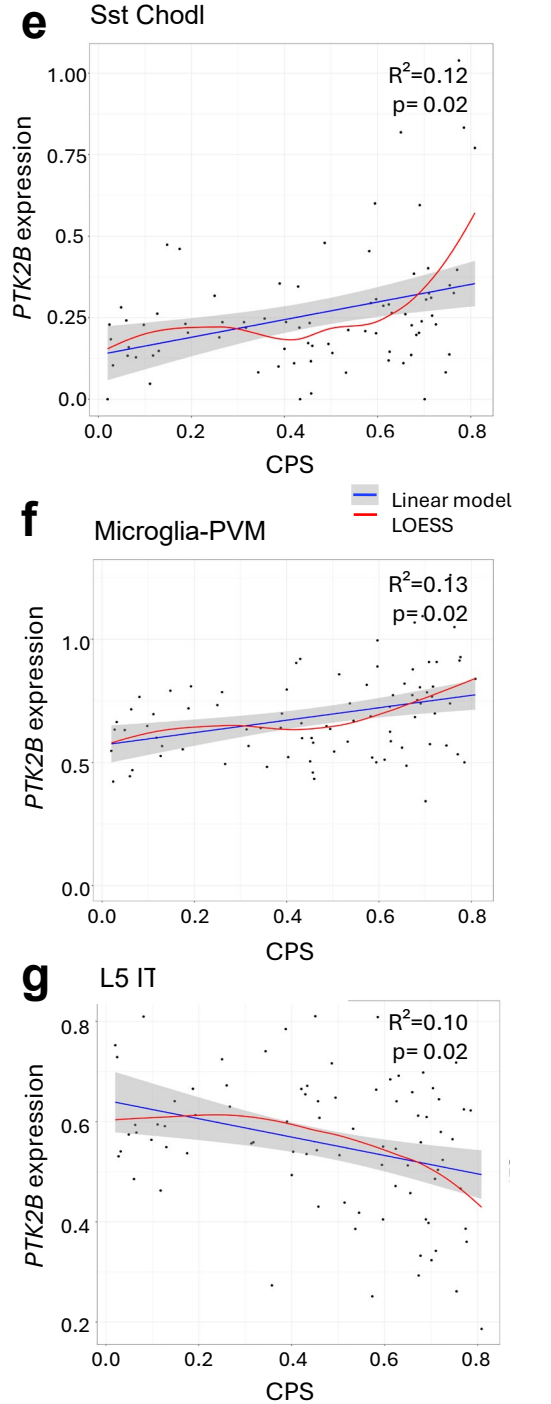

Supplementary Figure 2

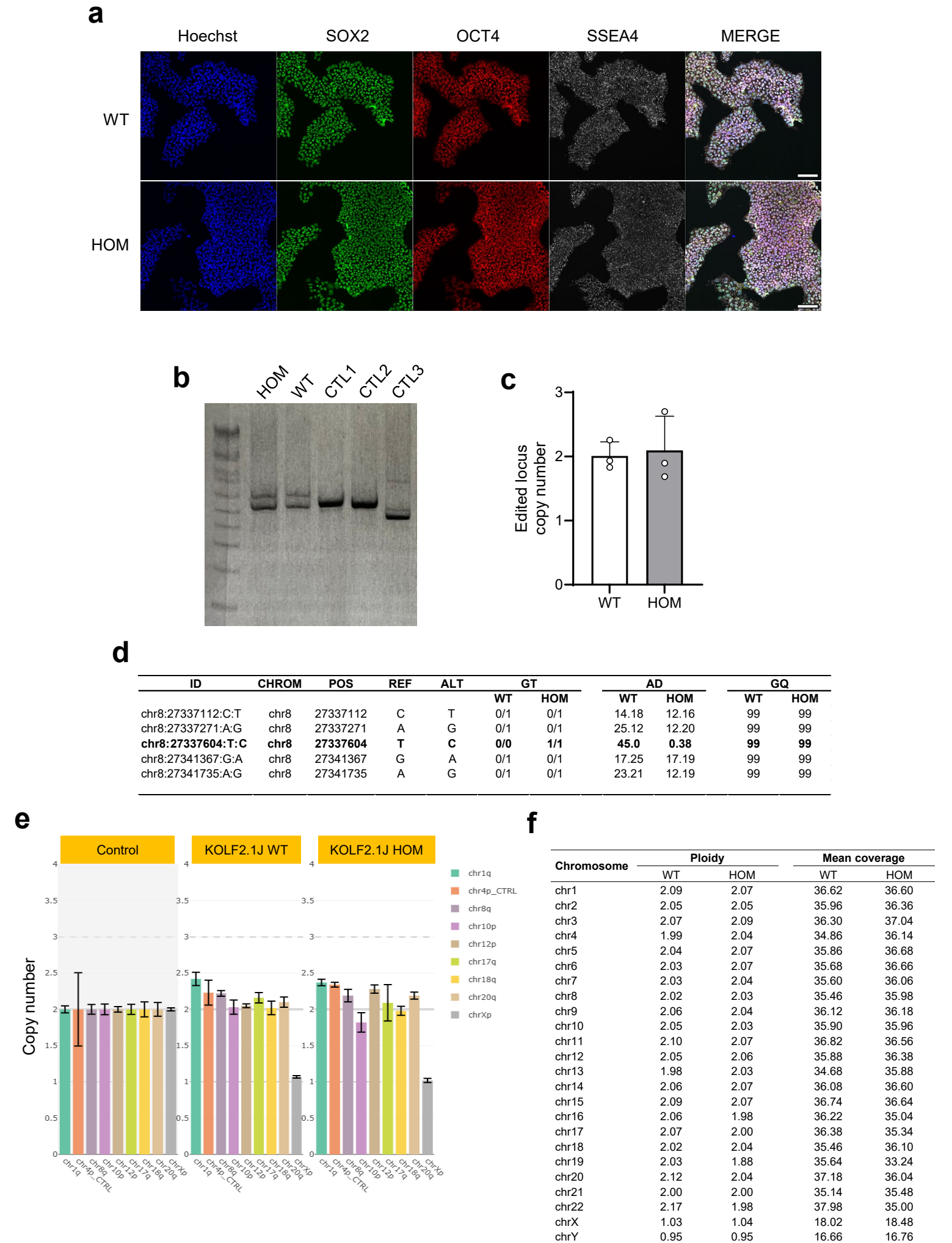

Supplementary Figure 3

**a**

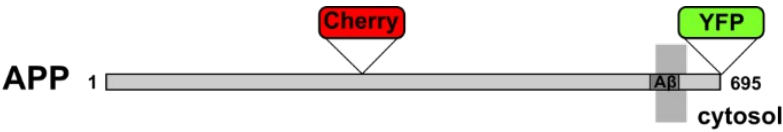

**b**

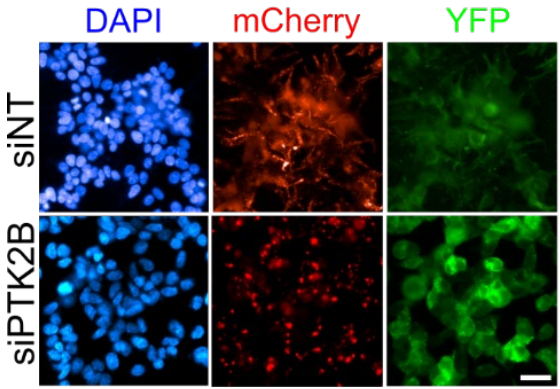

**c**

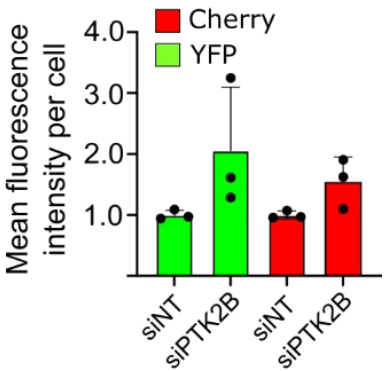
